# Improving Seedling Survival for Forest Restorations: A Novel Screening Method to Identify Microbial Allies Against Drought Stress

**DOI:** 10.1101/2025.04.24.650507

**Authors:** Sonja Magosch, Claudia Barrera, Adrian Bölz, Karin Pritsch, Michael Rothballer, J. Philipp Benz

## Abstract

Improving drought tolerance of tree seedlings by plant growth-promoting microorganisms (PGPMs) is a promising approach for nature-based forest restoration. Identifying suitable microorganisms requires a robust selection, including efficient *in planta* screenings. We sampled at two forest sites in southern Germany with drought legacies and within a dry period to enhance the probability of isolating drought-tolerant microbes. Metabarcoding of the resident soil community revealed a broad on-site diversity with the potential for diverse plant growth-promoting and stress-resistance traits. We isolated 1,292 bacteria and 59 fungi from fine roots of Norway spruce and European beech. 429 isolates were identified on genus level. The most abundant genera were *Paraburkholderia* (121) and *Bacillus* (43) in bacteria and *Penicillium* (8) and *Umbelopsis* (8) in fungi. Isolates were scored *in vitro* for abiotic stress tolerance and plant growth-promoting traits, revealing diverse plant growth-promoting abilities for 31 bacteria and a particularly high stress tolerance for 8 fungi. Importantly, an axenic 24-well-plate system was developed to test the most promising bacteria on spruce seedlings under drought. The system allowed direct comparison of inoculation effects on seedling growth and survival with or without drought application. Two strains improved survival and root length under drought, while six strains significantly promoted plant growth under well-watered conditions. This study represents one of the first larger scale screenings for PGPMs isolated from forest soils on tree seedlings under drought and may contribute to finding nature-based drought mitigation strategies.

## 1. Introduction

Throughout the last decades, drought events have enhanced in duration and frequency (Schlyter et al. 2006; Lobell, D. B. & Gourdji, S. M. 2012; Parmesan, C. & Hanley, M. E. 2015). They are characterized by below-average precipitation and high evaporation rates over a longer period, often accompanied by increased temperatures (Dai 2011). In European forests, increased drought results in elevated tree mortality (Bolte et al. 2016; Maguire, A. J. & Kobe, R. K. 2015), often leading to failed recovery or shifts in tree species composition (Hartmann et al. 2022). Forests account for one third of the terrestrial area and are of high relevance for ecosystems and humans (Bundesministerium für Ernährung und Landwirtschaft 2018). They constitute important players in climate regulation and protection by, for instance, nutrient cycling and storage of high amounts of anthropogenic and atmospheric carbon (Canadell, J. G. & Raupach, M. R. 2008; Hagedorn et al. 2016). As a result, shifts or degradation of forests become major issues for global carbon fluxes.

On a plant level, dry periods lead to restricted soil nutrient and water assimilation as well as morphological changes including reduced height and size of leaves, stems, shoots and roots (Koza et al. 2022; Bogati, K. & Walczak, M. 2022). Drought stress is often indicated by a reduced relative water content and transpiration rate, which are important cooling factors to diminish heat damage (Abdelaal et al. 2021). This is particularly relevant in seedlings (Sigala et al. 2020; Walters et al. 2023). Due to their shallow root system and growth in upper soil layers, they are more prone to desiccation by evapotranspiration, while water uptake during dry periods is restricted. Thus, seedlings are more susceptible to drought stress compared to adult trees (Goke A. & Martin P. H. 2022; Mediavilla, S. & Escudero, A. 2003). Moreover, in contrast to mature trees, seedlings do not possess enough carbon reserves that can be utilized during stress periods to sustain physiological functions (Goke A. & Martin P. H. 2022). In order to prevent water loss, stomata are closed early, leading to a lower O_2_-/CO_2_ exchange, carbon starvation (Domingo et al. 2023; Hartmann et al. 2022) and finally to the accumulation of harmful reactive oxygen species (Fadiji et al. 2022; Zhang et al. 2023). Moreover, biochemical processes like the regulation of endogenous phytohormone levels such as auxin or abscisic acid, which are involved in plant defense against stresses and plant growth regulation, are disturbed under drought (Egamberdieva et al. 2017; Ahluwalia et al. 2021). Forests in Central Europe are currently dominated by Norway spruce (*Picea abies*) and European beech (*Fagus sylvatica*), making up 30% of the forest area (Pretzsch et al. 2014). Compared to beech, spruce is even more susceptible to abiotic and biotic disturbances such as limited water supply, storms, and pathogen attacks, among other things due to its superficial root system and an associated reduced competitiveness (Pretzsch et al. 2014; Pretzsch, H. & Schütze, G. 2005; Schlyter et al. 2006; Han et al. 2022). In many parts of central Europe and Scandinavia, Norway spruce is of high commercial importance and is the preferred timber for construction (Perstorper et al. 1995; Huber et al. 2023; Ramage et al. 2017; Hartmann et al. 2022), highlighting the critical importance for preserving this tree species.

Restoration of forest sites relies on the successful establishment of seedlings, which is particularly challenging under drought (Sigala et al. 2020; Gaviria, J. & Engelbrecht, B. M. J. 2015). Different measures have been tested to mitigate the negative impacts of drought on seedling survival, growth, and stress resilience. One option is to use chemical fertilizers as they support faster growth (Wang et al. 2018; Adesemoye et al. 2009). However, this strategy is associated with high costs, can negatively affect the environment, and contributes to the depletion of non-renewable resources (Pahalvi et al.; Hakim et al. 2021; Fahde et al. 2023). Other, more sustainable options are recently gaining importance. The application of biofertilizers comprising plant beneficial bacteria and fungi has been a promising approach in agricultural production systems (Abdelaal et al. 2021), and inoculation with plant growth-promoting microorganisms (PGPMs) is an eco-friendly and financially advantageous alternative to chemical fertilizers (Fahde et al. 2023). PGPMs are free-living in the rhizosphere soil, rhizoplane, phyllosphere or endosphere, and can promote plant growth or reduce biotic or abiotic stresses through a wide range of functional traits (Fadiji et al. 2022; Cao et al. 2023; Compant et al. 2010) involving direct and indirect mechanisms (Finkel et al. 2020). Direct ways comprise promotion of plant growth by phytohormone production and improved nutrient supply, while indirect strategies include pathogen suppression and induced systemic resistance (Glick 2012; Fahde et al. 2023; Kuhl et al. 2021).

Microbial plant growth promotion involves the production of siderophores, 1-aminocyclopropane-1-carboxylate (ACC) deaminases, production of phytohormones like indole-3-acetic acid (IAA) or abscisic acid, phosphate solubilization and nitrogen fixation (Koza et al. 2022). Additionally, tree seedlings inoculated with microbiota that had previously experienced drought stress, showed a higher survival rate under drought (Allsup et al. 2023) making the application of stress-primed microorganisms an important strategy for mitigating the negative effects of drought.

Many plant growth-promoting bacteria (PGPB) and fungi have been characterized regarding their effectivity for agriculture, and some of those are already commercially utilized, e.g. the bacterial genera *Azospirillum* spp., *Bacillus* spp., *Burkholderia* spp., *Pseudomonas* spp. and *Streptomyces* spp., as well as *Serratia* spp. and *Rhizobium* (Fahde et al. 2023; Glick 2012; Abdelaal et al. 2021; Cao et al. 2023; Fadiji et al. 2022). Fungi being applied in agriculture comprise genera like *Trichoderma* spp., *Aspergillus* spp., or arbuscular mycorrhizal fungi (Zin, N. A. & Badaluddin, N. A. 2020; Nevalainen 2020). *P. abies* and *F. sylvatica* form ectomycorrhizae with specific fungal groups in the Basidio- and Ascomycota (Nickel et al. 2018). Their beneficial effects also in tree seedling establishment and their role in enhancing drought tolerance is being intensively studied (Lehto und Zwiazek 2011; Grams et al. 2021; Pretzsch et al. 2014; Nickel et al. 2018). However, their wider application is limited by the difficulty to cultivate most of them, and to establish the symbiosis in the field. In our study, we focused on other and more easily culturable PGPMs including bacteria and root-associated, mainly saprotrophic, fungi.

Despite the root microbiome of trees being of central importance for nutrient and water acquisition and protection against pathogens (Fracchia et al. 2021), there is only very limited information on PGPMs in boreal and temperate forest ecosystems (Puri et al. 2020a; Radhapriya et al. 2018). Li et al. 2022 isolated a plant growth-promoting *Paenibacillus* strain from a spruce forest in China, but its PGP potential was only shown for clover and *Arabidopsis*. There are some rare examples of PGPB tested for spruce (Lehr et al. 2008, Schrey et al. 2005) where seedlings were inoculated with the plant-beneficial *Streptomyces* AcH505 strain, which was found to protect the seedlings from *Heterobasidion annosum* infection, to work as mycorrhizal helper bacterium, promote root branching and slightly increase lateral root formation. Another study identified a *Pseudomonas* strain and two *Bacillus* strains with beneficial effects on hybrid spruce growth, and to increase seedling dry weight compared to uninoculated control plants (Chanway et al. 2000).

However, to date, no studies have investigated the use of PGPMs to improve drought tolerance in spruce seedlings. To support successful seedling establishment, a rapid and targeted screening for a broad range of PGPMs is essential, ideally involving a direct application to the plant of interest.

Aiming to overcome these limitations, a screening for plant-beneficial bacterial and fungal isolates was initiated that included, in addition to a classical *in vitro* screening, also a newly developed axenic plant cultivation system using 24-deep-well-plates with spruce seedlings, being particularly drought-sensitive. To increase the probability of finding species being effective under drought, isolates were extracted after one week of low or no precipitation from soils with moderate-dry to dry climate legacies in southern Germany. Prior to isolation, the on-site diversity was analyzed by amplicon sequencing to estimate the potential of obtaining a broad strain collection with PGP and stress resistance properties. By isolation and selection via *in vitro* assays, promising isolates were filtered out from the total microbial pool.

## 2. Material and methods

### 2.1 Study sites and soil sampling

Samples for isolation and amplicon sequencing were taken in Summer 2021 from two different mixed stands of Norway spruce (*Picea abies*) and European beech (*Fagus sylvatica*) in southern Germany. At the moderately dry site Kranzberger Forst (MAP 854 mm, MAT 8.2 °C (Wilhelm et al. 2023)), located at longitude 11°39’39.6’’E, latitude 48°25’8.4’’N, samples were taken in late July from plots previously subjected to drought-stress (Pretzsch et al. 2014). At the drier site Kelheim (MAP 713 mm, MAT 8.5 °C (Wilhelm et al. 2023)), located at longitude 11°49’19.2’’E, latitude 48°56’8.16’’N, samples were taken in the beginning of September. Samples were taken after 7 days of no or low precipitation (Table S1). Detailed information about the two sites can be found in Pretzsch et al. 2014.

For amplicon sequencing with the aim to analyze the microbial community composition at the time of sampling, at each site 5 x 5 soil cores (4.5 cm diameter to a depth of 10 cm) were taken from below 3 neighboring beech or spruce trees. Samples were immediately frozen, transported on dry ice and stored at -80 °C until usage.

For isolation of bacteria and fungi, fine root samples were taken directly below the trees by tracking them from coarser roots, with 5 samples from beech and spruce, respectively, at each site. Each fine root sample consisted of 3-5 fine roots (<1 mm diameter) with adhering soil. They were kept at 4 °C upon transport and processed on the day of sampling to avoid changes in the microbial composition.

### 2.2 Sample preparation for amplicon sequencing to characterize the soil microbial communities

DNA was extracted from roots with adhering soil using the FastDNA Spin kit for soil (MP biomedicals, Hessen, Germany), following manufacturer’s instructions. DNA concentration was measured at a Nanodrop® One spectrometer (Thermo Fisher Scientific, Waltham, MA, USA) and adjusted to 1.2-15 ng µL^-1^. DNA-samples were sent to the Core Facility of Microbiome, ZIEL, TUM (Freising, Weihenstephan; PD Dr. Klaus Neuhaus) for library preparation, purification, and Next Generation Sequencing (Illumina MiSeq® reagent kit v3 (600 cycle), paired-end reads). For sequencing of the v4 region of the 16S rRNA gene, the primers 515F (GTGYCAGCMGCCGCGGTAA, Parada et al. 2016) and 806R (GGACTACNVGGGTWTCTAAT, Apprill et al. 2015) were used. The fungal ITS2 region was amplified using a mixture of different primers according to Tedersoo et al. 2015 with adaptations: ITS3-Mix123-fw (AHCGATGAAGAACGCAG), ITS3-Mix45-fw (CATCGATGAAGAACGTRG), ITS4-Mix1-rv (TCCTCCGCTTATTGATATGC), ITS4-Mix2-rv (TCCTGCGCTTATTGATATGC), ITS4-Mix3-rv (TCCTCGCCTTATTGATATGC) and ITS4-Mix4-rv (TCCTCCGCTGAWTAATATGC). After demultiplexing, the data was provided from the Core Facility of Microbiome as fastq files. Sequencing data was processed using QIIME2 (v2024.2.0, (Bolyen et al. 2019) with the standard trimming and denoised using DADA2 (Callahan et al. 2016). Taxonomic classification of Amplicon sequence variants (ASVs) was performed via Greengenes2 (McDonald et al. 2024) for the 16S amplicons, while ITS amplicons were classified using the “feature-classifier” trained on the UNITE database (Abarenkov et al. 2024). Samples with less than 8000 reads, in the case of 16S, or less than 3000 reads, in the case of fungi, as well as singletons were removed. Overall, 95 samples were retained while 5 samples were removed due to a low sequencing depth. ASVs found in the non-template controls and in low abundance were removed. Samples were rarefied to the lowest depth using the “Rarefy” function of the GNuniFrac packages (Chen et al. 2012). Community composition was evaluated using the microbiome packages (v1.24.0, Leo et al. 2017). To compare the similarities between the different groups, the Bray-Curtis distance was calculated using the vegan package (v2.6.6, Oksanen et al. 2017), and a Principal Coordinates Analysis (PCoA) was performed. The functional profile was predicted using Tax4Fun2 (Wemheuer et al. 2020) with the default database and a 97% similarity cut-off. Then, the predicted enzymes were manually inspected to identify those involved in the pathways evaluated in the *in vitro* assays. All plots were produced using ggplot2 (Wickham 2016).

### 2.3 Bacterial and fungal isolation and cultivation

Fine root samples were grinded in 1 mL 1x phosphate buffered saline (1x PBS) using autoclaved mortar and pestle, diluted up to a factor of 10^-3^, and 100 µL of the suspension was plated on petri dishes on different solid agar media in 10 replicates per medium: King’B medium (Fluka Biochemika, Germany), M9 minimal medium (after Arie Gerloff), and MMNC medium (Kottke et al. 1987). For isolation of *Streptomyces*, 500 µL of the grinded soil samples were transferred into HNC liquid medium (Nonomura H. & Hayakawa M. 1988) and incubated for 30 min at 120 rpm and 42 °C. The suspension was filtered through glass wool and diluted up to a factor of 10^-1^ and 10^-2^ and 100 µL of both dilutions were plated on KM4- and HV-agar (Shirling, E. B. & Gottlieb, D. 1966; Hayakawa, M. & Nonomura, H. 1967). Plates were incubated for 1-2 weeks at 27 °C. For isolation of fungi, the same dilutions were used as for bacteria and 100 µL were plated on solid MMNC medium plates, and for some samples additionally on King’s B and M9 medium. Two fungal isolates (*Gymnopus* F32, *Lycoperdon* F31) were isolated by placing small pieces from inside the fruiting bodies on MMNC medium. All plates were incubated for up to 10 days at 25 °C. For purification and subculturing, part of the mycelium or spores were transferred to new MMNC medium where they were grown for 10 days at 25 °C and daily inspected for potential bacterial contamination.

Bacteria were pre-cultivated in NB liquid medium at 180 rpm and 28 °C for 2-3 days until they reached the stationary phase. They were diluted at a factor of 1:100 in fresh NB medium and cultivated overnight. These starting cultures were used for experiments in their exponential growth phase. Bacteria of the genus *Streptomyces* were cultured in baffled flasks to avoid cell clumping.

Fungi were cultivated on cellophane membranes (Einmachfolie 1-2-3, Deti GmbH, Germany) on MMNC agar. Since all fungi used in the *in vitro* assays were non-sporulating, they were inoculated with mycelial pieces of defined sizes (∼19.6 mm^2^). They were grown for 7-10 days at 25 °C in the dark. Mycelial pieces gained from parts of the actively growing mycelium were used for inoculation.

### 2.4 Sequencing of the 16S rRNA gene or ITS region for identification of the isolates

390 bacteria and 39 fungi were identified on a genus level, chosen based on different morphologies. For identification of bacteria, colony PCRs were performed. Due to the rigidity of *Streptomyces* colonies, DNA was in this case extracted from liquid cultures using the DNeasy UltraClean Microbial Kit (Biotechnologies, Madison, USA), following manufacturer’s instructions, and 20-50 ng µL^-1^ were used for PCR. The v1-9 region of the 16S rRNA gene was amplified with the primers 27F (5’-AGAGTTTGATCCTGGCTCAG-3’) and 1492R (5’- ACGGYTACCTTGTTACGACTT-3’) (Metabion International AG, Germany). For identification of fungi, DNA was extracted from 5-10 mg mycelium with the Animal and Fungi DNA Preparation Kit (Jena Bioscience, Germany) according to manufacturer’s instructions. The ITS region was amplified using around 100 ng µL^-1^ DNA and the primers ITS1-F (TCCGTAGGTGAACCTGCGG) and ITS4-R (TCCTCCGCTTATTGATATGC) (White 1990). PCR results were verified by agarose gel electrophoresis (1% agarose in 1x TAE buffer, 0.005% Midori Green, loading dye 3x OrangeG, 180 V, 37 min). The unpurified PCR products were sent for unidirectional sequencing (Eurofins genomics, Germany, seq2plate kit). The FASTA sequences were blasted using NCBI (https://blast.ncbi.nlm.nih.gov/Blast.cgi). Sequences with >97% identity were assigned to the respective genera.

### 2.5 Identification of potentially beneficial isolates based on different in vitro assays

36 bacterial and 17 fungal isolates were screened for their plant growth-promoting and abiotic stress resistance abilities in different *in vitro* assays, regardless of their origin (tree species/site). Due to the high number of isolates, the selection of bacteria and fungi for characterization and screening within *in vitro* assays was based on the 16S and ITS rRNA sequencing results and literature research. Genera, which were already known to contain plant growth-promoters, specifically in spruce, were preferably, albeit not exclusively, selected. Bacterial starting cultures were prepared as described above. If not stated otherwise, cells were harvested by centrifugation and washed twice in 1x PBS before each assay. Fungi were grown on MMNC agar plates as described above using pieces from the actively growing mycelium for inoculation during the assays. All *in vitro* assays were performed in three replicates. Based on the performance of the isolates in the different assays, scores were assigned with higher numbers standing for a better performance in the assays. The highest score to be reached for each *in vitro* assay was 3. For bacteria, which were subjected to 3 stress tolerance assays and 5 plant growth promotion assays, the possible maximum score was 24. For fungi, the possible maximum score was 12, based on 2 stress tolerance (osmotic stress and salt stress) and 2 plant growth promotion assays (phosphate solubilization, IAA-production).

#### Siderophore production

For the detection of siderophore production, the CAS-overlay agar method of Pérez-Miranda et al. 2007 and Lynne et al. 2011 was used with modifications. After cell washing, the optical density was measured at 600 nm (OD_600_) and adjusted to 0.5. 50 µL of each bacterial suspension was spotted on NB agar plates and incubated at 28 °C. After 48 h, 10 mL of the CAS-overlay agar (Chrome Azurol Blue S (Sigma Aldrich, USA), FeCl_3_ (Fluka Biochemika, Germany), HDTMA (Sigma Aldrich, USA) mixed 1:10 with 32.24 g L^-1^ Piperazin-N’N-bis-(2-ethanesulfonic acid) in 0.9% agar, pH 6.8) was pipetted on top of the bacterial colonies. Siderophore production was visible after 12 h by a color change from blue to orange.

#### 1-Aminocyclopropane-1-carboxylate (ACC) deaminase production

The enzyme 1-aminocyclopropane-1-carboxylate (ACC) deaminase stimulates plant growth by hydrolyzing the ethylene precursor ACC (Glick 2012). Bacterial growth on minimal medium with ACC as single nitrogen source indicates the presence of an ACC deaminase. The assay was performed according to Brown, C. M. & Dilworth, M. J. 1975 and Rothballer et al. 2008. The OD_600_ of the bacterial suspensions was adjusted to 0.5. 25 µL were spotted on M9 medium with 9.35 mM NH_4_Cl, 3 mM ACC (Thermo Fisher Scientific, Waltham, MA, USA) or no nitrogen source. After incubation at 28 °C for 10 days, the utilization of ACC as nitrogen source was determined by comparing bacterial growth on the three different media. The strain *Variovorax* sp. M92526_27 (Kuhl et al. 2021) was used as an ACC-utilizing positive control.

#### Identification of potentially nitrogen-fixing bacteria

For identification of nitrogen-fixing bacteria, two different media were used. Nitrogen-free semi-solid Nfb-agar was prepared according to Döbereiner 1995. Cell suspensions were adjusted to an OD_600_ of 0.1 and 10 µL were spotted on top of 5 mL Nfb-agar into glass tubes. After incubation for 144 h, fixation of atmospheric nitrogen was visible by formation of pellicles inside of the agar. Bacterial single colonies were additionally streaked out on nitrogen-free Jensen’s agar (in g L^-1^: sucrose, 20; K_2_HPO_4_, 1; MgSO_4_ · 5 H_2_O, 0.5; NaCl, 0.5; FeSO_4_ · 7 H_2_O, 0.183; Na_2_MoO_4_, 0.005; CaCO_3_, 2) according to Kuhl et al. 2021. Colony growth indicates fixation of nitrogen. The nitrogen fixation assay was only considered positive if bacteria were able to grow on both media. *Azospirillum brasilense* Sp7 was used as a positive control.

#### Phosphate solubilization

Phosphate solubilization was tested on National Botanical Research Institute’s phosphate (NBRIP) growth medium (in g L^-1^: glucose, 10; Ca_3_(PO_4_)_2_, 5; MgCl_2_ · 6 H_2_O, 5; MgSO_4_ · 7 H_2_O, 0.25; KCl, 0.2; (NH_4_)_2_SO_4_, 0.1; pH 7.0) according to Nautiyal 1999. For bacteria, the OD_600_ was adjusted to 0.5 and 25 µL were spotted on NBRIP agar plates. After incubation at 28 °C for 10 days, phosphate solubilization was visible by formation of a clear zone around the bacterial spots. Phosphate solubilizing *Luteibacter* sp. Cha2324a_16 (Kuhl et al. 2021) was used as a positive control. For fungi, sterile cellophane membranes were added on the NBRIP agar surface and inoculated with mycelial pieces. Plates were incubated for 10 days at 25 °C in the dark. For evaluation, the cellophane membranes were removed making the clear halos visible. Some halos were clearer and some only lightly indicated compared to others, being responsible for the different scores (0-3) assigned in case of fungi.

#### Production of indole-3-acetic-acid

The production of indole-3-acetic acid (IAA) was determined using the colorimetric method of Gordon, S. A. & Weber, R. P. 1951. Bacterial pre-cultures were prepared as described above and transferred into NB liquid medium with and without 1.5 mg mL^-1^ of the IAA-precursor L-tryptophane (Carl Roth, Germany). They were incubated for 72 h at 28 °C and 180 rpm in the dark. The OD_600_ was adjusted to a unified value (lowest OD_600_ measured). Cells were harvested at 5000 xg for 2 min and 100 µL of the supernatant was mixed with 100 µL of Salkowski reagent (0.01 M FeCl_3_ anhydrous in 35% perchloric acid) and 1 µL orthophosphoric acid in a 96 well plate. After 40 min of incubation, the IAA content was measured at 530 nm in a plate reader (SpectraMax iD3, Molecular Devices). For the preparation of a standard curve, commercial indole-3-acetic acid (Fluka Biochemika, Germany) at concentrations from 0-100 µg mL^-1^ was used. For data evaluation, the OD_600_ was normalized to 1. *Herbaspirillum frisingense* GSF30 served as a positive control. IAA amounts >5 µg mL^-1^ were considered as positive IAA production. For fungi, 3 mL MMNC liquid medium with and without 1.5 mg mL^-1^ L-tryptophane were inoculated with 2 mycelial pieces in 24-deep-well-plates. After incubation for 96 h, 100 µL of the supernatant were mixed with 100 µL of the Salkowski reagent and 1 µL orthophosphoric acid in 96 -well-plates and proceeded as described above. For evaluation, IAA production was normalized to the fungal dry biomass. Values above a threshold value of 0.5 µg mg^-1^ were considered as positive IAA production.

#### Tolerance to different abiotic stresses

To examine their tolerance to different abiotic stresses (salt, osmotic, pH), 2 µL of the bacterial main cultures were transferred into 200 µL NB liquid medium with seven different concentrations of either NaCl, or PEG, and H^+^ (Table S4). For salt stress induction, NaCl was used (modified from Carvalho et al. 2014) and to measure osmotic stress tolerance, different amounts of polyethylene glycol 6000 (PEG, Serva Electrophoresis GmbH, Germany) were applied. The amount of PEG was based on decreasing water potentials using the formula of Michel, B. E. & Kaufmann, M. R. 1973, according to Kumar et al. 2014 and Jayakumar et al. 2020, with 326 g L^-1^ PEG creating an osmotic pressure of -1.2 MPa. With a pH value of around 5, forest soils are relatively acidic (Bundesministerium für Ernährung und Landwirtschaft 2018). Therefore, the bacterial tolerance to low pH values was also assessed. The tolerance to different pH values was analyzed by adjusting the pH of NB liquid medium using NaOH and HCl (modified from Kuhl et al. 2021). All assays were performed in sterile 96-well-plates and the OD_600_ was measured after 0, 24 and 48 h. Bacterial recovery from the 2-3 highest concentrations of NaCl, PEG and H^+^ was determined by plating 25 µL of these concentrations on NB agar plates, incubating them at 28 °C for 48 h, and observing bacterial growth. Fungi were subjected to salt and osmotic stress. The tolerance to low pH values was not tested, since most fungi grow at a pH range of 5 to 6 (Basu et al. 2015), matching the demands for growth on forest soils. NaCl tolerance was tested on agar plates containing MMNC medium with seven different NaCl concentrations (Table S4). They were grown for 10 days at 25 °C in the dark. The agar surface area covered by fungal mycelium was measured for each plate using ImageJ. PEG tolerance was assessed in liquid, adding 2 mycelial pieces to 3 mL MMNC medium with seven different PEG concentrations (Table S4). They were incubated for 96 h at 130 rpm and 25 °C. The mycelium was harvested and washed thoroughly 2x in MilliQ water. Mycelia were dried at 65 °C and the dry weight was measured.

### 2.6 Application of 29 bacterial isolates to Picea abies seedlings in a newly developed 24-well-plate format

An axenic 24-well test system was developed to examine the effect of bacteria characterized within the *in vitro* assays on the growth of *P. abies* seedlings (Figure 1). To set up the system, 24-deep-well-plates were filled with custom-made substrate (TS1 fein with white peat (0-5 mm), perlite (1-7.5 mm) and black peat, 3:1:1 (v/v), pH 5.5, without NPK-fertilizer; Klasmann-Deilmann GmbH, Germany). In case of well-watered conditions, 3.25 mL ddH_2_O and for drought conditions, 0.75 mL ddH_2_O were added to all wells. After autoclaving, 0.5 mL ½ MMN pH 5.5 liquid medium without carbon source was added to each well. *P. abies* seeds were obtained from the Bavarian State Forests Nursery (Bayerische Staatsforsten AöR, Laufen, Germany). They were washed in ddH_2_O for 30 min and sterilized for 45 min in 35% H_2_O_2_, rinsed with ddH_2_O for 30 min and placed onto MMNC square petri dishes for germination for 1 week in the dark in a tilted position. To inoculate the seedlings, bacterial main cultures were washed twice in 1x PBS, and the cell density was adjusted to 10^8^ cells per mL. The seedlings were incubated in the bacterial suspension for 1 h at 28 °C and 100 rpm. Each inoculated seedling was transferred into one well of the prepared 24-well-plates. The lower part of a phytatray (Sigma-Aldrich, Munich, Germany) was used as a translucent cover for the 24-well-plates. To enable gas exchange, small holes were burned into the lid using a hot needle. The lid was fixed with parafilm on top of the plates. The 24-well test systems were kept in a phytochamber under controlled conditions (8/16 day/night cycle, 55% humidity, 20 °C, 25 Wm^-2^ light intensity). After 3 weeks, plant morphology was evaluated by measuring root and shoot fresh weight as well as length, and seedling dry weight after drying the seedlings for 3 weeks at room temperature. Moreover, bacterial rhizosphere competence was estimated by counting colony forming units (CFUs) after 3 weeks of plant growth in a phytochamber. To this end, three roots per treatment (inoculated/non-inoculated, drought-stressed/well-watered) were grinded in a sterilized mortar in 1x PBS and diluted up to a factor of 10^-4^. 100 µL of the 10^-3^ and 10^-4^ dilution were plated on NB medium, and the CFUs were counted after incubation of 24-48 h at 28 °C. Only plants with vital root tips were chosen for evaluation.

**Figure 1:**
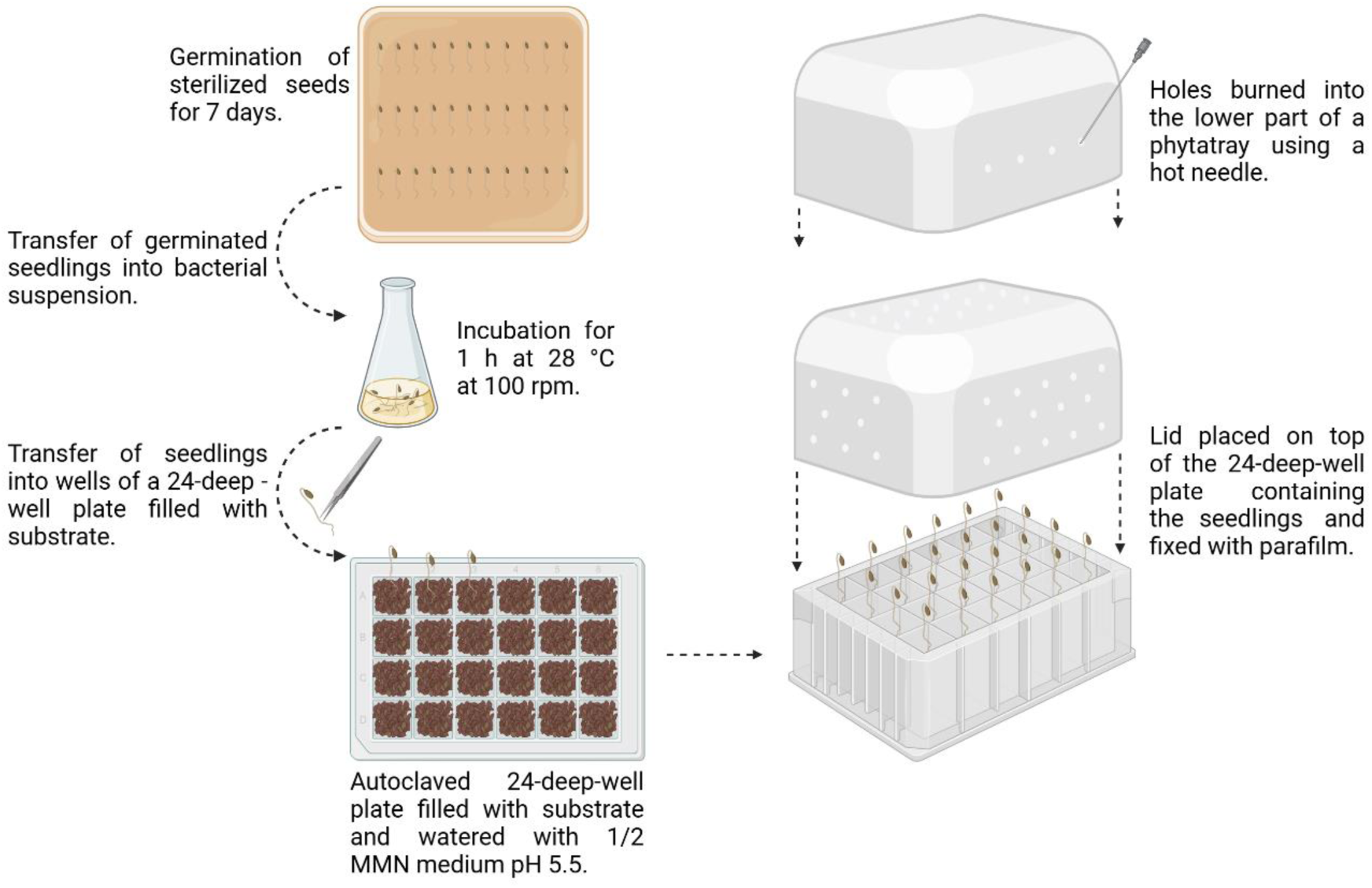
Newly developed test system using 24-deep-well-plates. *P. abies* seeds are germinating for 1 week on MMNC agar plates before they are inoculated in bacterial suspensions or 1x PBS (control) for 1 h. They are transferred into 24-deep-well-plates filled with sterilized soil. After burning holes into the lower part of a phytatray with a hot needle, plates are sealed with the modified lid and put into a phytochamber for 3 weeks. This figure was created with BioRender.com.

In the initial experiments using the 24-well test system, internal controls were included on each plate to monitor potential variances between the plates. Six control plants and twelve inoculated plants were used, with one row of six wells left empty between the two treatments to prevent potential cross-contamination. No variance between the control plants was observed during the plant experiments. Consequently, to increase the number of replicates, the internal controls were omitted, and the control plants placed in a separate plate, allowing for 24 replicates for both the control and inoculated plants. It was demonstrated that the internal controls remained consistent across different plates (Figure S4).

### 2.7 Statistical analysis

Origin Pro (version 2021b) was used for preparing figures (except for microbiome data). Data of the plant parameters were tested for normal distribution by Shapiro-Wilk test. In case normal distribution was rejected, non-parametric Mann Whitney test was performed. Data was analyzed with Fisher’s least significant test and one-way ANOVA (p-value 0.05). Details regarding the statistics for amplicon sequencing are provided in the corresponding section.

## 3. Results

### 3.1 The resident root-associated microbial community at both sites and tree species bears the potential for isolating PGP and stress resistant microbes

Given that both chosen sampling sites (Kranzberg and Kelheim) are (moderately) dry, environmentally distinct and host two different tree species, the resident root-associated microbial community at the time of sampling after one week of low or no precipitation (Table S1) was analyzed by amplicon sequencing to visualize the existing diversity and evaluate its potential for isolating PGP and stress-resisting candidates. On both sites, a total of 3364 Amplicon sequence variants (ASVs) were found for bacteria, and 4357 ASVs for fungi. The resident root-associated microbial community was highly similar across both sites and tree species, exhibiting only minor differences. The fungal community showed a separation by tree species as well as by location (Figure S2 B), while differences in the bacterial community composition were influenced more by tree species than by sampling site (Figure S2 A). The major difference between sites was a higher relative abundance of Bacillota in Kranzberg ASVs, which was even more pronounced for spruce compared to beech (Figure S2 C). In case of fungi, Mortierellomycota were highly represented in spruce in Kranzberg. In Kelheim, a higher abundance of Basidiomycota was detected compared to Kranzberg (Figure S1 A).

A total of 30 bacterial phyla were identified in the soil microbiome (Figure 2 A). The top 10 most abundant phyla comprised Pseudomonadota, Acidobacteriota, Actinomycetota, Verrucomicrobiota, Bacteroidota, Bacillota, Cyanobacteriota, Chloroflexota, Thermoproteota and Desulfobacterota (in descending order). Overall, 15 different phyla were found in the fungal microbiome (Figure S1 A). Ascomycota was the dominant phylum, followed by Basidiomycota. Mortierellomycota was the third most represented phylum.

**Figure 2:**
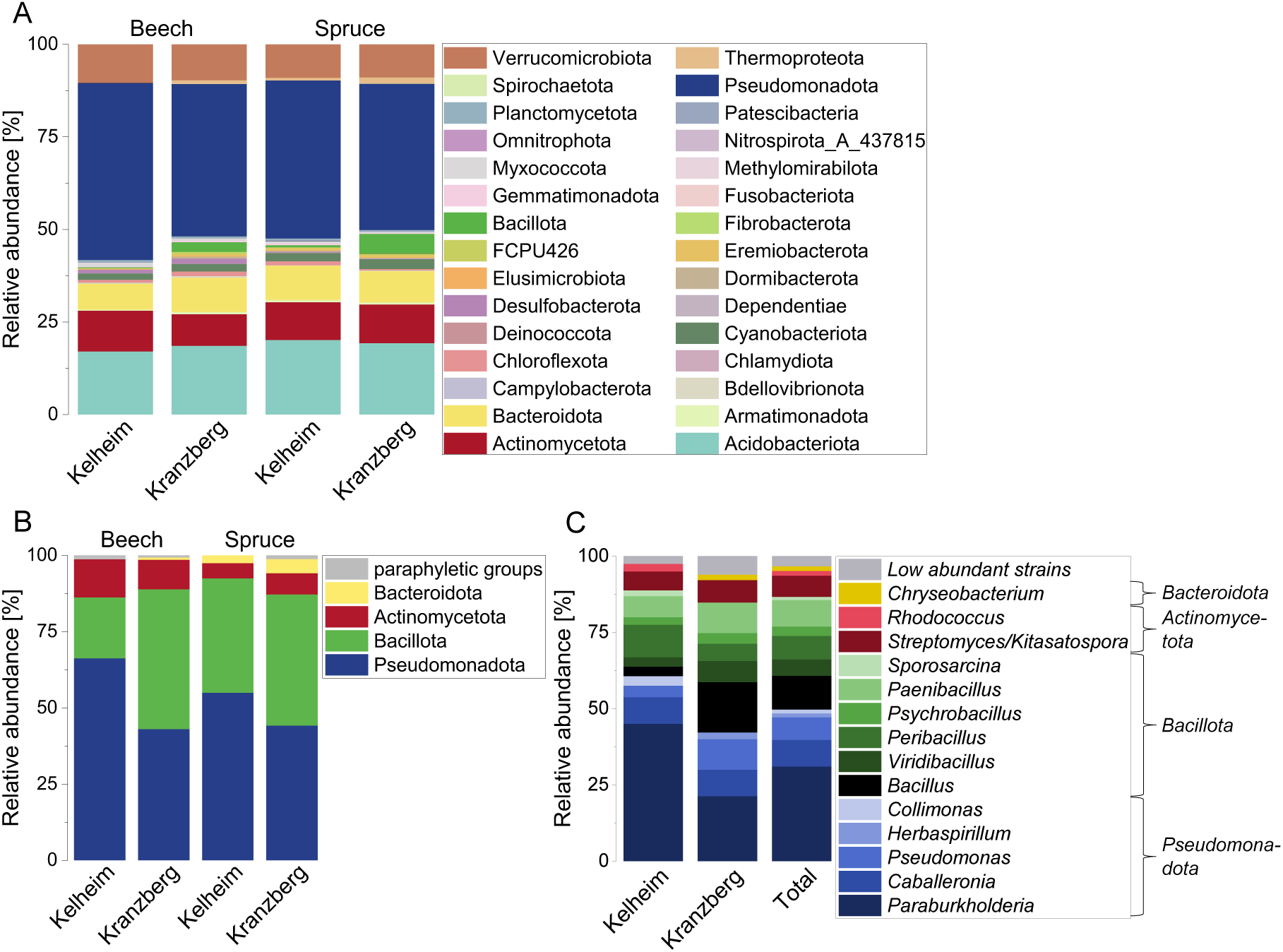
Comparison of the relative abundance of bacterial phyla in rhizosphere communities and root isolates of beech and spruce in Kelheim and Kranzberg. (A) Relative abundance [%] of the bacterial phyla in beech and spruce rhizosphere communities in Kelheim and Kranzberg. (B) Relative abundance [%] of the bacterial phyla isolated as single strains from beech and spruce roots in Kelheim and Kranzberg. (C) Relative abundance [%] of the bacterial genera isolated as single strains in Kelheim and Kranzberg, and at both sites combined. The designation “Low abundant strains” refers to genera with less than 3 isolates. Bacterial genera of the same phylum are depicted in similar colors. Pseudomonadota: blue. Bacillota: green. Actinomycetota: red. Bacteroidota: yellow. More closely related genera are depicted in similar colors.

The bacterial amplicon sequencing data was further subjected to a functional prediction analysis to provide an initial characterization of the microbial community and to assess its potential for identifying PGP and stress resistant isolates (Figure 3). The trait with the highest predicted abundance was IAA-production followed by N-fixation. The potential to harbor bacteria with the desired traits (PGP and stress resistance) was similar across both sites and tree species.

**Figure 3:**
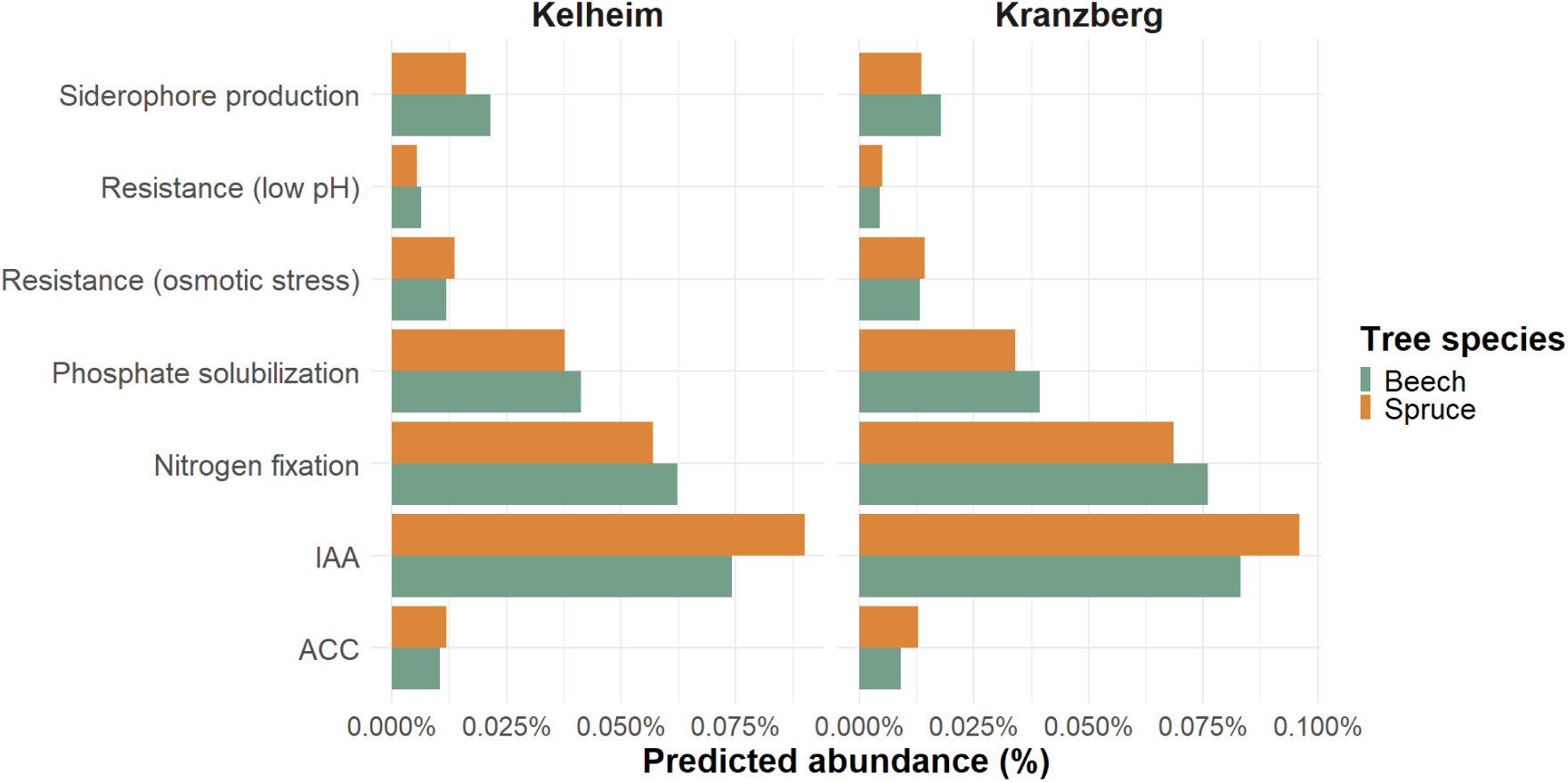
Trait prediction of the community analysis. Predicted abundance [%] of different enzymes for the bacterial beech and spruce rhizosphere community in Kelheim and Kranzberg being involved in the PGP and stress resistance pathways evaluated in the *in vitro* assays of this study.

### 3.2 Isolation and identification of soil microorganisms

All in all, 1,292 bacteria and 59 fungal isolates were obtained from soils in Kranzberg (moderate-dry site) and Kelheim (dry site). Based on their morphology on agar plates, bacteria were assigned to about 20 morphotypes. From each morphotype, between 10-20 strains were subjected to Sanger sequencing, leading to the identification of 390 bacteria, which could be assigned to 25 different genera. Overall, 230 bacteria originated from Kranzberg (23 different genera) and 160 from Kelheim (15 different genera). The bacterial isolates belonged to four different phyla (50.5% Pseudomonadota, 38.2% Bacillota, 8.7% Actinomycetota, 1.8% Bacteroidota) with a low proportion assigned to paraphyletic groups (0.8%) (Figure 2 B). The most frequently isolated genus was *Paraburkholderia* with 121 isolates (31.0%), followed by *Bacillus* with 43 isolates (11%), and *Caballeronia* and *Paenibacillus* with 34 isolates (8.7%) (Figure 2 C). *Bacillus* was isolated more often in Kranzberg compared to Kelheim (16.5% vs. 3.1%). In Kelheim, *Paraburkholderia* accounted for 45% of all genera, while in Kranzberg, a more even distribution of the different bacterial genera was observed. In addition to bacteria, 39 fungi were identified on genus level, with 25 isolated from Kelheim and 14 from Kranzberg. Their genus-level identification revealed 14 different genera (13 in Kelheim, 4 in Kranzberg) belonging to 3 different phyla (56.4% Ascomycota, 38.5% Mucoromycota, 5.1% Basidiomycota) (Figure S1 B). *Penicillium* and *Umbelopsis* were the most frequently isolated fungal genera with 20.5% each, followed by *Mortierella* with 12.8% (Figure S1 C).

### 3.3 Screening of microbial isolates for their tolerance to abiotic stresses and plant growth-promoting activities

Since no substantial differences were observed in the microbial rhizosphere communities between sites or species (neither in terms of relative abundance nor functional prediction), the isolates were screened for their plant growth-promoting and abiotic stress resistance abilities in different *in vitro* assays, regardless of their origin (beech/spruce, Kelheim/Kranzberg). In total, 36 bacterial strains were characterized, with 20 isolated from Kelheim and 16 from Kranzberg. Additionally, 17 fungal strains were chosen for characterization, with 13 from Kelheim and 4 from Kranzberg (Table S2 and Table S3). The choice of isolates for further characterization was guided by literature research, while trying to incorporate a diverse set of genera.

### 3.4 Bacteria showed a high PGP potential in vitro

Abiotic stress tolerance assays included different concentrations of NaCl, PEG, or different pH values and each tested strain was evaluated with a scoring system based on the test outcome (methods). *Streptomyces* Ke434, *Rhodococcus* Ke466 and *Streptomyces* KF215 showed the highest tolerance, reaching a score of 7 out of 9 in the stress tolerance assays (Table 1; Figure 5 A). *Streptomyces* Ke434 exhibited tolerance to an osmotic potential of -1.5 MPa, whereas *Rhodococcus* Ke466 and *Streptomyces* KF215 tolerated up to -1.25 MPa. The average PEG tolerance for bacteria was -0.78 ± 0.24 MPa. The highest NaCl concentration was tolerated by two *Rhodococcus* strains (Ke442, Ke466), two *Streptomyces* strains (Ke434, KF143) and *Sporosarcina* Ke477, withstanding concentrations of up to 5.5% (Table 1), while the average NaCl tolerance was 2.14 ± 1.96%. Almost all strains were able to grow at a pH of 5. Eight strains even grew at pH 4, including six *Paraburkholderia* strains (Ke15, Ke24, Ke162, Ke296, Ke341, Ke398), *Rhizobium* Ke26 and *Caballeronia* Ke57 (Table 1). The average pH tolerance was 4.79 ± 0.74. None of the strains was able to grow below a pH of 4. Most of the bacteria were also able to recover from the 3 highest stress levels (Table S4 and Table S6) Recovery from low pH values seemed to be most difficult, since only a few isolates could recover from pH values below 4, namely *Bacillus* Ke157, *Rhodococcus* Ke466 and *Viridibacillus* KF108, which recovered after incubation in NB medium with a pH of 2.

**Table 1:**
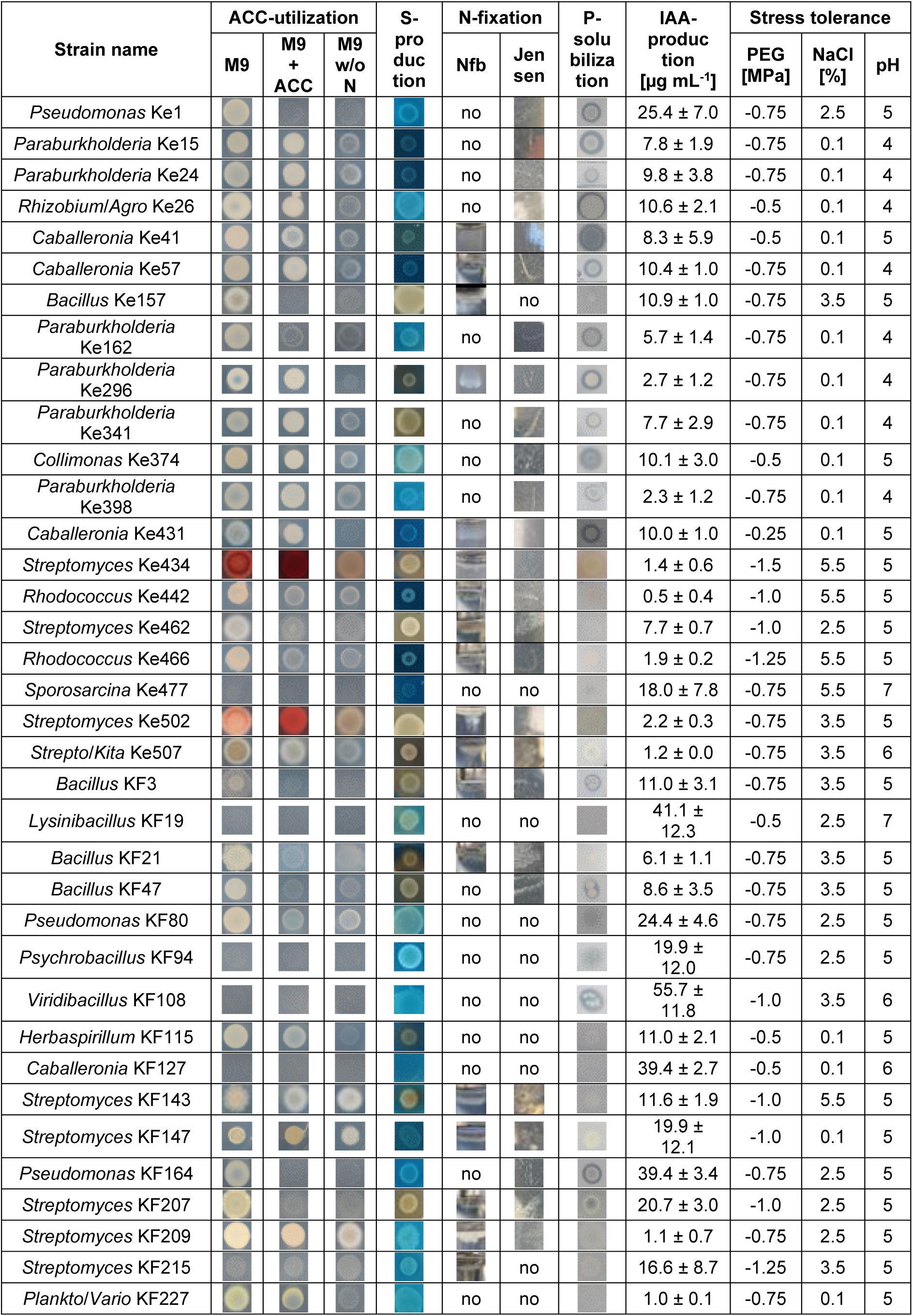
Summary of all bacterial *in vitro* assays. , comprising ACC-utilization, siderophore production, N-fixation, P-solubilization, IAA-production and tolerance to osmotic stress, salt stress and low pH-values.

In addition to stress tolerance, the plant growth-promoting (PGP) potential was also evaluated by assessing the bacteria’s ability to solubilize phosphate, produce siderophores, fix nitrogen, utilize ACC, and produce IAA. *Caballeronia* Ke41, *Paraburkholderia* Ke296 and *Streptomyces* KF207 showed the highest plant growth-promoting abilities with scores of 13 and 12 out of 15 (Table 1; Figure 5 A). 16 of 36 strains (∼ 44%) were able to produce siderophores, particularly from the genera *Bacillus* and *Streptomyces*. Solubilization of phosphate was detected in 19 of 36 strains (∼ 53%) and included all *Paraburkholderia* strains. ACC-utilization was observed in 20 strains (∼ 56%), including all *Paraburkholderia* and most *Streptomyces* strains. Regarding nitrogen fixation, 16 of 36 strains (∼ 44%), especially represented by *Rhodococcus* and *Streptomyces*, showed growth in both tested nitrogen-free media, indicating their nitrogen-fixing potential. The positive controls *Variovorax* sp. M92526_27 (ACC utilization), *Azospirillum brasilense* Sp7 (nitrogen fixation) and *Luteibacter* sp. Cha2324a_16 (phosphate solubilization) also tested positive in their respective assays (Table S5). All bacteria were also examined for their ability to produce the phytohormone indole-3-acetic acid (IAA). 28 out of the 36 strains (78%) produced IAA at concentrations above the threshold level of 5 µg mL^-1^ (Table 1). 11 strains produced IAA concentrations above 15 µg mL^-1^ (score of 3) with *Viridibacillus* KF108, *Lysinibacillus* KF 19, *Caballeronia* KF127, and *Pseudomonas* KF164 reaching the highest concentrations of 55.7, 41.1, and 39.4 µg mL^-1^ (Table 1). The positive control *Herbaspirillum frisingense* GSF30 produced 12.4 ± 2.1 µg mL^-1^ IAA. Some of the tested strains even showed an IAA-production above the threshold level of 5 µg mL^-1^ after growth without tryptophane (Table S7).

Overall, 31 out of 36 strains (86%) exhibited PGP abilities in at least two different *in vitro* assays, while only five strains showed positive results in just one PGP assay (Table 1; Figure 5 A). The combined results of the bacterial *in vitro* assays were subjected to a Principal Component Analysis (PCA) to identify potentially genus-specific commonalities showing as clusters (Figure 4). A cluster mainly comprising the genera *Paraburkholderia* and *Caballeronia* was represented by strong osmotic stress tolerance (PEG), as well as P-solubilization and ACC-utilization abilities. Actinomycete isolates of the genera *Streptomyces* and *Rhodococcus* clustered by comparably strong N-fixation, siderophore production and salt stress tolerance abilities. Since the isolated genera were not equally distributed between the two different sites, a statement about site-related effects is not possible.

**Figure 4:**
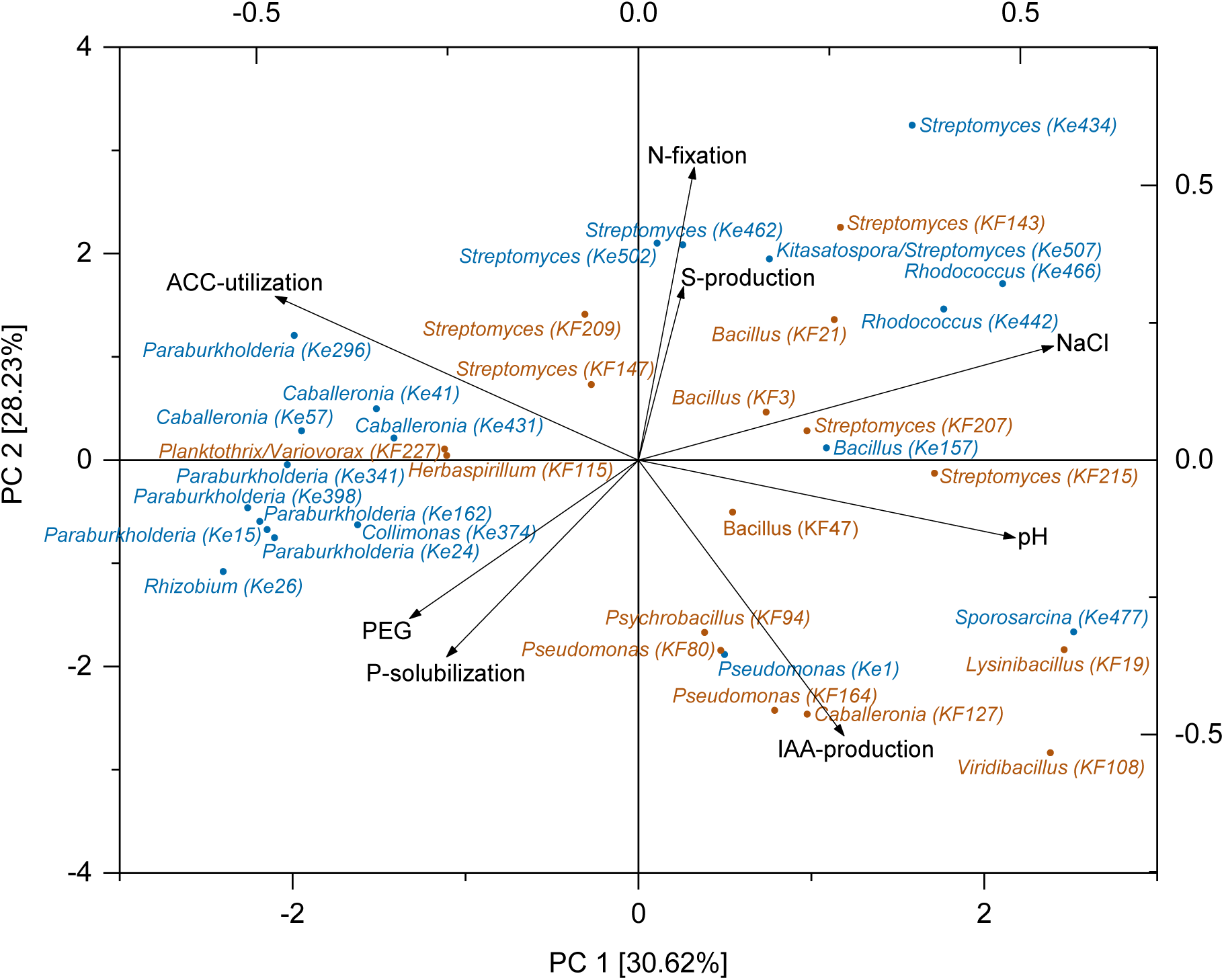
Principal Component Analysis (PCA) identifying genus related clusters for the *in vitro* assays of the bacterial strain collection. Variables: pH, PEG and NaCl tolerance, as well as production of IAA based on measured values; production of siderophores, phosphate solubilization, ACC-utilization and nitrogen fixation based on the presence/absence of the PGP trait (1/0). Isolates from Kelheim are marked in blue and isolates from Kranzberg in brown.

*Streptomyces*, *Paraburkholderia*, *Caballeronia*, *Collimonas*, *Rhodococcus* and *Bacillus* ranked among the ten best performing bacterial genera of the *in vitro* assays. Except for *Collimonas* and *Rhodococcus*, the best performers belonged to the most abundant genera of the strain collection, highlighting a high prevalence of PGPB among the isolates.

### 3.5 Fungi tolerated especially high NaCl and PEG concentrations

Fungal stress tolerance at different NaCl and PEG concentrations was generally high, with 8 out of 17 strains (47%) reaching the maximum score of 6 (Figure 5 B). The average osmotic tolerance was at -1.13 ± 0.49 MPa. *Metapochonia* F6 and three *Umbelopsis* strains (F11, F12, F14) grew well even at the highest PEG concentration tested (-1.75 MPa) (Table 2).

**Figure 5:**
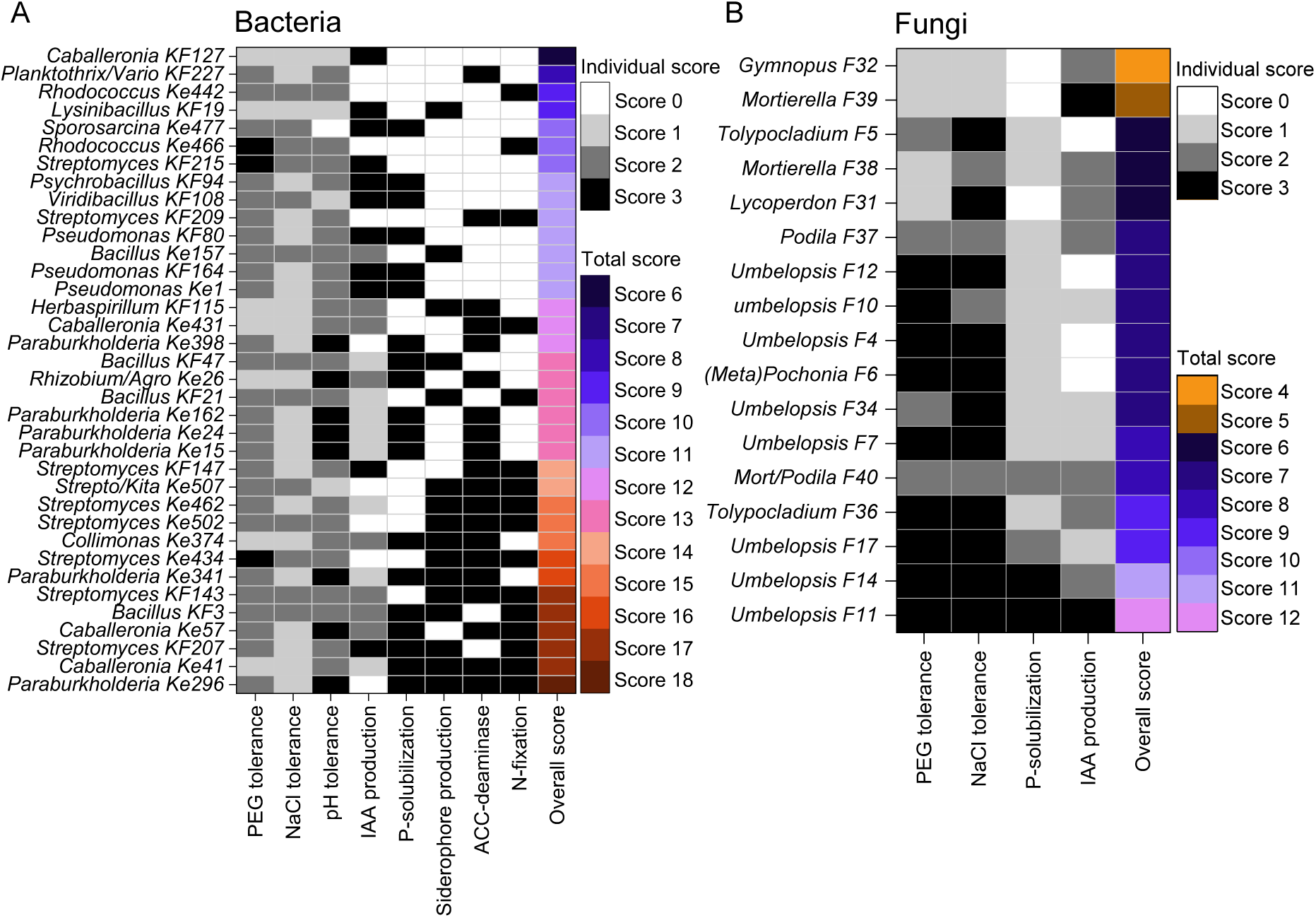
Heatmaps showing the scoring of the isolates in the different *in vitro* assays. The isolates are sorted by ascending overall scores, given in the column on the right. (A) For bacteria PEG-, NaCl- and pH-tolerance, as well as IAA-production were evaluated with scores ranging from 0 3. P-solubilization, siderophore-production, ACC-utilization and N-fixation were either evaluated with 0 for “not detectable” or 3 for “detectable”. (B) For fungi PEG- and NaCl-tolerance, as well as P-solubilization and IAA-production were evaluated with scores ranging from 0-3.

**Table 2:** Summary of all fungal *in vitro* assays. , comprising P-solubilization, IAA-production, and tolerance to osmotic stress and salt stress. Some halos observed in the P-solubilization assay were clearer and others only lightly indicated, being responsible for the different scores assigned (0-3).

| Strain | P-solubilization | | IAA-production<br>[$\mu\text{g mg}^{-1}$] | Stress tolerance assays | |
| --- | --- | --- | --- | --- | --- |
|  | Reaction on medium | Evaluation |  | PEG [MPa] | NaCl [%] |
| *Umbelopsis* F4 | | + | $0.36 \pm 0.15$ | -1.5 | 7.5 |
| *Tolypocladium* F5 | | + | $0.50 \pm 0.2$ | -0.75 | 7.5 |
| *Metapochonia/Pochonia* F6 | | + | $0.05 \pm 0.03$ | -1,75 | 12 |
| *Umbelopsis* F7 | | + | $0.59 \pm 0.38$ | -1.25 | 7.5 |
| *Umbelopsis* F10 | | + | $0.89 \pm 0.08$ | -1.5 | 5.5 |
| *Umbelopsis* F11 | | +++ | $5.53 \pm 2.25$ | -1,75 | 7.5 |
| *Umbelopsis* F12 | | + | $0.43 \pm 0.15$ | -1,75 | 7.5 |
| *Umbelopsis* F14 | | +++ | $1.75 \pm 0.42$ | -1,75 | 7.5 |
| *Umbelopsis* F17 | | ++ | $0.82 \pm 0.34$ | -1.25 | 7.5 |
| Fruiting body, *Lycoperdon* F31 | | - | $1.74 \pm 0.09$ | -0.5 | 7.5 |
| Fruiting body, *Gymnopus* F32 | | - | $1.78 \pm 1.07$ | -0.5 | 0.1 |
| *Umbelopsis* F34 | | + | $0.90 \pm 0.27$ | -1.0 | 7.5 |
| *Tolypocladium* F36 | | + | $1.41 \pm 0.23$ | -1.25 | 7.5 |
| *Podila* F37 | | + | $2.21 \pm 0.37$ | -1 | 3.5 |
| *Mortierella* F38 | | + | $4.73 \pm 1.71$ | -0.5 | 3.5 |
| *Mortierella* F39 | | - | $6.35 \pm 1.46$ | -0.5 | 0.1 |
| *Mortierella/Podila* F40 | | ++ | $3.99 \pm 1.29$ | -0.75 | 3.5 |

*Metapochonia* F6 showed a high dry weight (DW) at all PEG concentrations, with a maximum DW of 37.4 ± 6.6 mg at -1.0 MPa. Since several other *Umbelopsis* strains, such as F10, F11, F12, F14 and F17, had similarly elevated dry weights, this might indicate a potential usage of PEG as a carbon source by these strains. The average NaCl tolerance was at 6.07 ± 3.03%. *Metapochonia* F6 exhibited the highest detectable salt tolerance, growing at 12% NaCl, followed by ten strains that grew at up to 7.5% NaCl (Table 2).

Regarding their PGP potential, almost all fungi solubilized phosphate (14 of 17, ∼ 82%), although to different extents (Table 2), and 13 out of 17 fungi (∼ 77%) produced IAA above the threshold value of 0.5 µg mg^-1^ (Table 2). Two fungi showed an especially high IAA-production: *Umbelopsis* F11 with 5.5 ± 2.3 µg mg^-1^ and *Mortierella* F39 with 6.4 ± 1.5 µg mg^-1^, both reaching the maximum score of 3 for IAA-production. In contrast to bacteria, fungi were not able to produce IAA without tryptophane.

Among the bacterial isolates, *Paraburkholderia* Ke296 reached the highest overall score (18), followed by two *Caballeronia* strains (Ke41 and Ke57), two *Streptomyces* strains (KF147 and KF207) and *Bacillus* KF3, each with a score of 17 (Figure 5 A). Two fungal isolates, *Umbelopsis* F11 and *Umbelopsis* F14 reached the highest scores of 12 and 11, respectively (Figure 5 B).

### 3.6 Application of bacteria to a newly developed 24-well-plate test system reveals two bacteria promoting spruce growth under drought

For further testing of pre-selected isolates to promote seedling drought tolerance, a 24-well-plate system was developed for a fast and axenic screening *in planta* (Figure 1 and methods). Pre-experiments showed neglectable variation between plates, and controls for cross-contamination were consistently negative. The final plate design allowed to test each selected isolate on 24 individual seedlings under control and drought conditions, respectively. Re-isolation 3 weeks after inoculation revealed colonization rates ranging from 4.9 x 10^3^ to 7.7 x 10^7^ colonies per mg root except for *Psychrobacillus* KF94 and *Viridibacillus* KF108, which could not be re-isolated (Table S10). 29 of the 36 bacterial isolates were tested in the 24-well system. Several showed significant positive effects on seedling growth under well-watered (WW) conditions (Table S9). However, two isolates clearly stood out due to their exceptionally pronounced PGP effects under drought stress (DS): *Caballeronia* Ke431 and *Paraburkholderia* Ke296. Both significantly increased seedling length mainly by an increase in root length under DS conditions compared to the uninoculated controls (Figure 6 A and B). Both strains were even able to increase root length compared to the WW control (Figure S11). Strikingly, *Caballeronia* Ke431 increased seedling survival 3-fold and *Paraburkholderia* Ke296 maintained survival (1.2-fold increase), both under DS (Table S10). Moreover, although *Psychrobacillus* KF94 and *Viridibacillus* KF108 could not be re-isolated from spruce roots, seedling survival increased about 3-fold after inoculation with the two isolates, and KF94 also significantly increased seedling dry weight under DS, but had no other beneficial effects on plant growth (Table S10, Figure S10 B).

**Figure 6:**
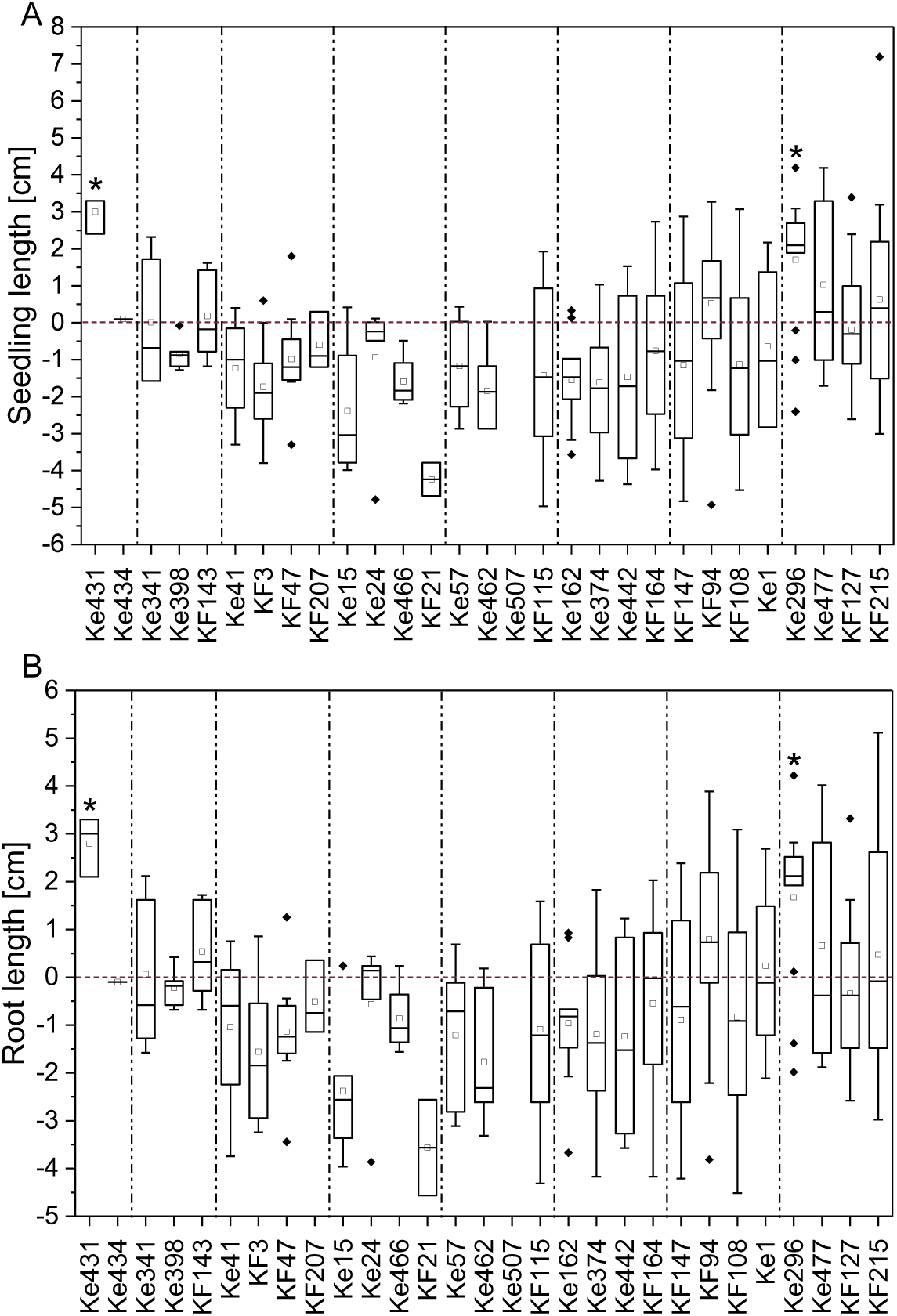
Seedling growth performance under drought. Increase or decrease in (A) seedling length and (B) root length under drought stress conditions after inoculation with the bacterial isolates in comparison to the control plants of each experiment (red zero baseline). The single experiments are separated by dashed lines. Each positive value indicates an absolute increase in seedling length or root length of the inoculated plants compared to the respective controls, while negative values mark a decrease. Significant plant growth-promotion after inoculation with the isolates compared to the respective control is marked with asterisks.

*Caballeronia* Ke431 additionally significantly improved seedling and root length as well as seedling, root and shoot fresh weight under WW conditions (Figure 7, Figure S5, Figure S7 A, Figure S9 A).

**Figure 7:**
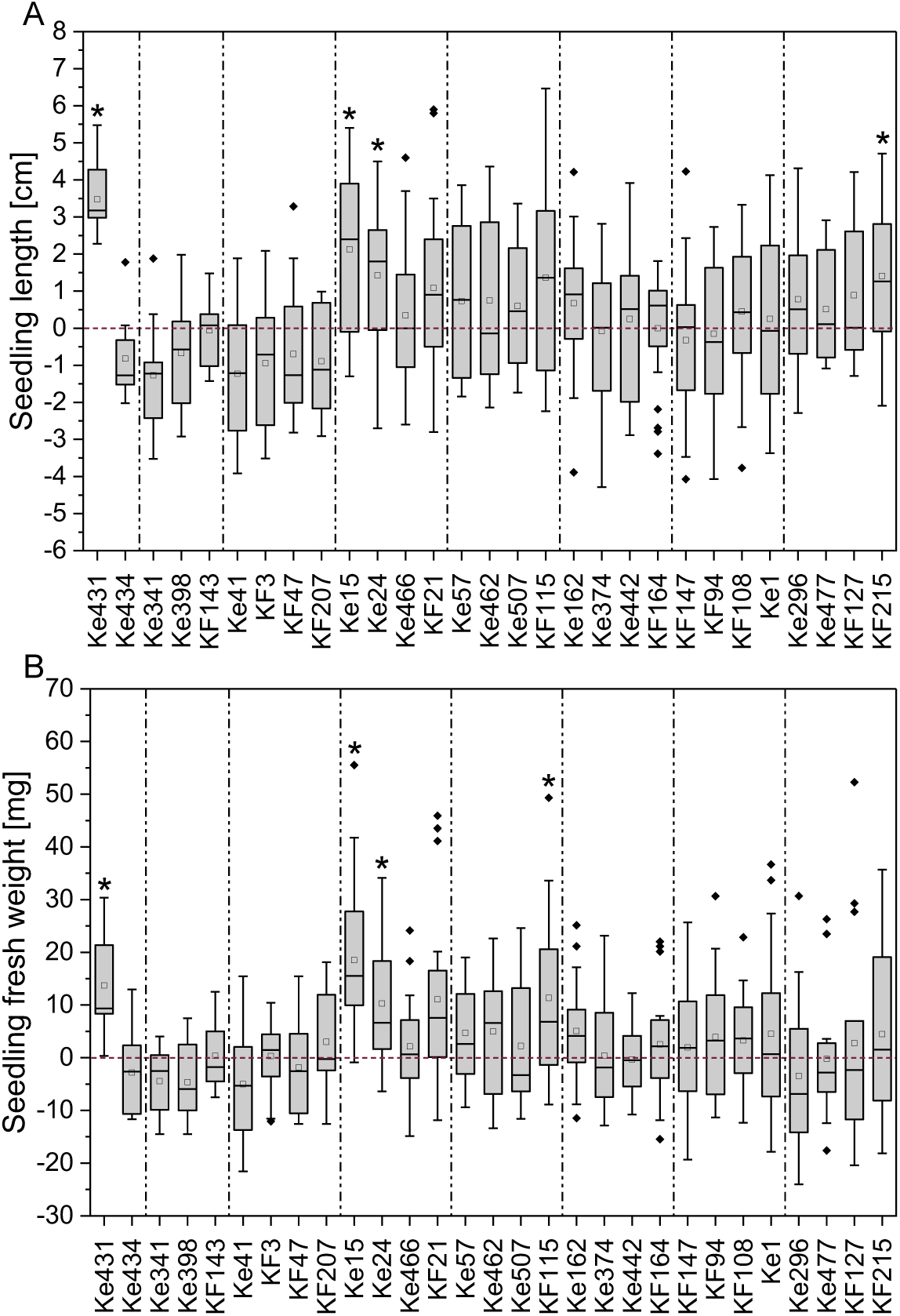
Seedling performance under well-watered conditions. Increase or decrease in (A) seedling length and (B) seedling fresh weight under well-watered conditions after inoculation with the bacterial isolates in comparison to the control plants of each experiment (red zero baseline). The single experiments are separated by dashed lines. Each positive value indicates an absolute increase in seedling length or seedling fresh weight of the inoculated plants compared to the respective controls, while negative values mark a decrease. Significant plant growth-promotion after inoculation with the isolates compared to the respective control is marked with asterisks.

Under WW conditions, also several other strains showed significant seedling growth-promoting effects (Table S9). Inoculation with *Paraburkholderia* Ke15 and Ke24, significantly increased seedling and root length, seedling, root and shoot fresh and seedling dry weight (Figure 7, Figure S5, Figure S7 A, Figure S9 A, Figure S10 A). Inoculation with *Streptomyces* KF215 significantly enhanced seedling length, root length and root fresh weight (Figure 7 A, Figure S5 and Figure S7 A). *Paraburkholderia* Ke162 inoculation increased root fresh weight (Figure S7 A) and *Herbaspirillum* KF115 showed significantly increased seedling, root and shoot fresh weight as well as dry weight (Figure 7 B, Figure S7 A, Figure S9 A and Figure S10 A). Inoculation with *Rhodococcus* Ke466 and *Bacillus* KF21 significantly increased seedling dry weight (Figure S10 A).

## 4. Discussion

### 4.1 The resident root-associated community bears a substantial potential for stress-resistant and plant growth-promoting microbes

The main aim of the present study was to isolate and identify bacteria and fungi with the potential to promote tree seedling growth under drought stress. To explore the available population diversity for isolation and to estimate the trait potential in the soil, a phylogenetic and functional characterization of the microbial rhizosphere community was performed. To capture a broad spectrum and increase the probability of isolating drought-adapted microorganisms, sampling was conducted at two different (moderately) dry sites from the rhizosphere of two tree species (spruce and beech) after a period with low or no precipitation (Table S1), as microorganisms can rapidly adapt to changes in environmental conditions (Gonzalez, J. M. & Aranda, B. 2023). Amplicon sequencing combined with a functional prediction analysis revealed a rhizosphere microbiome rich in taxa with known plant growth-promoting (PGP) capabilities, with only minor compositional differences observed between the two sites and tree species. The bacterial phyla Actinomycetota and Pseudomonadota, for instance, include many PGP and stress-resistant genera, such as *Streptomyces, Rhodococcus, Caballeronia* or *Paraburkholderia*. Their established role as stress-resistant PGPB in spruce and other plants (Glick 2012; Lehr et al. 2008; Schrey et al. 2005; Puri et al. 2020a, Puri et al. 2020b) positions these genera as promising candidates for the objectives of this study, and representatives of all these were also successfully isolated.

As anticipated, the amplicon sequencing data represented a much higher diversity compared to the strain collection. Nevertheless, all four bacterial phyla identified in the collection (Bacteroidota, Actinomycetota, Bacillota, and Pseudomonadota) were also among the top 10 most abundant phyla in the soil microbiome data (Figure 2 A and B). The genera *Paraburkholderia* and *Caballeronia* (both Pseudomonadota), which were isolated in high frequency, were also among the 8 most abundant genera in the bacterial rhizosphere community (Figure S3).

The bacterial diversity at both mixed spruce-beech study sites based on 16S rDNA amplicons matches previously observed phyla distributions found at other spruce and beech sites, being dominated by Acidobacteria (now Acidobacteriota), Proteobacteria (now Pseudomonadota), Actinobacteria (now Actinomycetota), Verrucomicrobia (now Verrucomicrobiota), Bacteroidota, Firmicutes (now Bacillota), and Chloroflexi (now Chloroflexota) (Uroz et al. 2016, Colin et al. 2017 Lladó et al. 2016 Choma et al. 2021, Haas et al. 2018 Tahovská et al. 2020). A high abundance of *Paraburkholderia* and *Caballeronia* in the present study confirms the presence of bacteria in the paraphyletic group *Burkholderia* (*sensu latu*) in the beech and spruce microbiome as observed in several studies (Uroz et al. 2016; Uroz, S. & Oger, P. 2017). As expected, the differences between the rhizosphere community and the strain collection were even more pronounced for fungi (Figure S1 A and B). The observed fungal diversity was nevertheless equivalent to other studies on spruce and beech rhizosphere microbiomes, being dominated by Basidiomycota, followed by Ascomycota and Mucoromycota (Choma et al. 2021; Haas et al. 2018; Uroz et al. 2016). On a genus level, the fungal microbiome is generally dominated by ectomycorrhizal genera such as *Cenococcum*, *Russula*, *Piloderma*, *Cortinarius*, *Hygrophorus* and (*Pseudo*)*Tomentella,* and also includes genera like *Archaeorhizomyces*, *Helotiales, Meliniomyces*, and *Penicillium* (Pretzsch et al. 2014; Nickel et al. 2018; Haas et al. 2018), which were all part of the 20 most abundant genera of the fungal microbiome observed in this study (Figure S3 B). The main fungal isolates in the present study were saprotrophic genera such as *Mortierella* and *Umbelopsis*, which were also reported in forest soils (Choma et al. 2021, Tahovská et al. 2020). The soil-dilution isolation method used in this study targets mainly root-associated saprotrophic fungi. Thus, an isolation of genera being part of the ectomycorrhizal group of fungi was not to be expected, and complex isolation methods would be required to increase the number of these isolates (Thorn et al. 1996).

### 4.2 Microorganisms with high stress tolerance and plant growth-promoting abilities were identified in a classical in vitro screening

While isolates and amplicon data were initially obtained separately for each site and tree species, all samples were combined for the identification of plant growth-promoting (PGP) microbes. Given the minor differences in the microbial community observed between sites and tree species, the focus was placed on the overall microbial community, enabling a comprehensive screening *in vitro*.

The strain collection of the present study exhibited multiple traits related to plant growth promotion, with 86% showing PGP abilities in more than one *in vitro* assay. *Paraburkholderia* Ke296 reached the highest overall score and was tested positive in all PGP assays except for IAA production, highlighting the high potential of this strain regarding plant growth promotion. When looking at the entire collection, however, IAA production was the most prevalent PGP trait, as already predicted by the functional analysis. The IAA levels produced in pure culture can serve as an indicator of a strain’s potential to mitigate the effects of osmotic stress on plant growth (Nadeem et al. 2016). Almost half of all bacteria produced IAA above the threshold value of 5 µg mL^-1^ even in the absence of tryptophane (47%). Although the tryptophane-dependent pathway is the most common in bacteria, especially gram-positive bacteria use alternative biosynthetic routes (Idris et al. 2007), which could thus partly be confirmed in the present study. Among the fungal isolates, 76% were also capable of producing IAA, including the genera *Umbelopsis* and *Mortierella*, which are already known for their ability to produce phytohormones such as IAA (Ozimek, E. & Hanaka, A. 2021; Yung et al. 2021; Ozimek et al. 2018).

The fungal isolates stood out particularly for their high stress tolerance with 47% of the strains growing on medium with high PEG (-1.25, -1.5, -1.75 MPa) and NaCl concentrations (7.5, 12, 15%) represented by several *Umbelopsis* strains, *Tolypocladium* and *Metapochonia* (Table 2). So far, little is known about the salt tolerance of the fungi studied here. *Umbelopsis* strains isolated from pine were able to tolerate 3% NaCl, although their growth rate was reduced (Min et al. 2014) and a *Tolypocladium* strain showed optimal spore production at up to 2% NaCl (Gardner & Pillai 1986). In comparison, the fungi tested in the present study showed a higher average NaCl tolerance of up to 6%. Moreover, some fungi of the present study showed even better growth on medium containing high compared to low PEG concentrations suggesting they were able to biodegrade PEG and use it as a carbon source as known for several microorganisms (Ayeni et al. 2022).

Members of the phyla Actinomycetota (*Streptomyces*, *Rhodococcus*) and Bacillota (*Viridibacillus*, *Bacillus* and *Sporosarcina*) showed the highest PEG and NaCl tolerance of the bacterial isolates. The high stress resistance observed for members of these phyla might be related to their ability to produce stress-resistant spores (Rao et al. 2022; Tovar-Rojo et al. 2003). *Streptomyces* Ke434 exhibited the highest PEG and NaCl tolerance (-1.5 MPa and 5.5%), making this strain an especially interesting candidate for seedling protection against osmotic stress. Several studies have demonstrated that *Streptomyces* and *Rhodococcus* strains can grow at NaCl concentrations ranging from 2.6-5.5%, with a few rare examples capable of growing in concentrations as high as 10% NaCl (Sadeghi et al. 2014; Palaniyandi et al. 2014; Carvalho et al. 2014). The tolerated NaCl concentrations are remarkably high for bacteria, as concentrations around 3.5-5% NaCl are typically associated with slightly to moderately halophilic bacteria (Oren 2008). The highest PEG concentrations tested in this study approached the threshold for severe osmotic stress, as plants begin to wilt irreversibly at a soil water potential of -1.5 MPa (Kuhl-Nagel et al. 2022). However, Bacillota and Actinomycetota are known to produce osmoprotectants, enabling them to tolerate high osmotic stress (Oren 2008; Ahluwalia et al. 2021).

Several PGP and stress resistance traits act synergistically with each other and work best in combination. Drought stressed plants inoculated with PGPB showed improved root and shoot growth and reduced cell damage that was connected with high IAA and ACC deaminase concentrations (Ahluwalia et al. 2021). ACC deaminase producing *Achromobacter piechaudii* ARV8, lowered ethylene levels in tomato plants and prevented plant growth inhibition under high salt and drought conditions (Glick et al. 2007). After inoculation with *Streptomyces* strains, beneficial effects on plant growth under NaCl and water stress were observed, such as production of high IAA and ACC deaminase levels being both correlated to abiotic stress alleviation in plants (Rao et al. 2022).

### 4.3 Double screening in form of a parallel in vitro and in planta screening increases the chances to isolate PGPMs

As a next and important step after the classical characterization of the isolates via different *in vitro* assays, a direct test for efficiency towards *P. abies* (spruce) seedlings under drought (as well as well-watered) conditions was implemented.

The efficacy of inoculation with bacteria depends on their rhizosphere competence for the specific host plant (Martínez-Vivero et al. 2010). 27 of 29 bacteria were successfully re-isolated from spruce roots after 3 weeks, demonstrating their rhizosphere competence and persistence at the root level. Only *Psychrobacillus* KF94 and *Viridibacillus* KF108 could not be re-isolated. The fact that their inoculation nevertheless increased plant survival suggests that they had a positive effect at least initially, but were unable to establish themselves at the roots. This phenomenon is not new. For example, *Azospirillum lipoferum* stimulated root growth in maize but could not be detected by quantitative PCR around 4 weeks after inoculation (Florio et al. 2017).

From the 86% isolates showing PGP to several degrees finally 2 out of 29 tested isolates supported seedling growth under drought. This underscores the limitations of *in vitro* screenings for identifying PGPMs, particularly in translating results from artificial conditions to real field environments (Abbamondi et al. 2016; Soumare et al. 2021; Smyth et al. 2011; Gamalero et al. 2022; Finkel et al. 2020) and the importance of suitable *in vivo* experiments. The fact that *in vitro* screening results are not directly transferrable to more natural *in vivo* conditions was true also for our study, in which all strains with beneficial effects *in vivo* exhibited only moderately high scores in the *in vitro* assays, ranging from 10 to 13 (except for *Paraburkholderia* Ke296, which achieved the highest recorded score of 18). *In vivo* experiments under more natural conditions are therefore essential for validating the efficacy of PGPM before proceeding with time-intensive field or greenhouse trials (Peng et al. 2023). The novel 24-deep-well-plate-based system was developed as a means to cover this gap between *in vitro* screenings and field application, enabling a controlled and fast screening of promising isolates on seedling fitness. In this approach, bacteria were applied directly to the target plant (spruce), avoiding the use of common model plants such as *Arabidopsis* or tomato, often used as proxies. The drought conditions applied in this system were severe, as indicated by a drop in seedling survival from over 75% (WW) to approximately 50% or lower under drought (Table S10).

The *in planta* screening revealed that two strains, *Caballeronia* Ke431 and *Paraburkholderia* Ke296, were able to promote spruce growth under DS, making them particularly noteworthy. These two strains also belonged to the eight most abundant genera of the bacterial rhizosphere community (Figure S3 A). As mentioned before, plants are often able to maintain their above-ground growth under moderate drought. Severe drought, on the contrary, leads to a decrease in shoot growth, whereas the roots continue to grow (Ahluwalia et al. 2021). This was also observed in the present study for spruce seedlings under drought, as sometimes control plants under DS showed an increased root length compared to control plants under WW conditions (Figure S11 B), although they had a reduced survival rate (Table S10). After inoculation with *Caballeronia* Ke431 and *Paraburkholderia* Ke296 under DS, this effect was especially pronounced (Figure S11). This would enable seedlings to reach deeper soil layers where more water is present while the upper soil layers are more affected by drought (Nickel et al. 2018; Walters et al. 2023). Incidentally, both DS-alleviating strains were able to fix nitrogen, for which there is evidence that this trait is involved in drought resistance mechanisms in seedlings: In *P. abies* for example, organic N-supply improved survival after planting in the field (Sigala et al. 2020).

Additionally, *Caballeronia* Ke431 as well as five other strains promoted spruce growth under WW conditions: *Herbaspirillum* KF115, *Streptomyces* KF215 and three *Paraburkholderia* strains (Ke15, Ke24, Ke162). Generally, root fresh weight was particularly positively affected by inoculation with the bacteria tested in this study, which may have resulted from their ability to produce IAA. It is well known that PGPB present in the rhizosphere or endosphere that are able to convert plant-produced tryptophane to IAA, can significantly promote plant growth (Puri et al. 2020a). Interestingly, IAA and ACC deaminase production were exhibited by six of the seven most-promising PGPB (Table S11), further indicating that these features could be particularly important for PGP. *Streptomyces* KF215 was not able to utilize ACC but showed a high tolerance to the osmotic stresses tested. All strains tolerated pH values between 4 and 5, being a prerequisite for surviving the relatively acidic pH of 5 in forest soils. Siderophore production and nitrogen fixation were the least common PGP traits, observed in only two strains each. The four *Paraburkholderia* strains shared most of their PGP abilities, indicating that some PGP traits might be genus-specific (Table S11).

Strains from all four PGP genera identified within the 24-well-based test system have already been shown to be plant beneficial in other studies. The genus *Herbaspirillum* has not been tested on spruce so far, but demonstrated PGP effects under DS for example in maize, where inoculation with *Herbaspirillum seropedicae* ZAE94 increased the yield up to 34% (Alves et al. 2015). An inoculation of hybrid white spruce with six bacteria, among them *Caballeronia sordidicola* LS-Sr, enhanced seedling growth by accumulating 607% more biomass in comparison to the control plants (Puri et al. 2020a). *Paraburkholderia phytofirmans* LP-R1r, *Caballeronia sordidicola* HP-S1r and *Caballeronia udeis* LP-R2r significantly increased spruce and pine seedling length by up to 60% and seedling biomass by up to 302% (Puri et al. 2020b). The plant beneficial *Streptomyces* AcH505 strain promoted root branching and lateral root formation in Norway spruce (Lehr et al. 2008, Schrey et al. 2005). However, these examples only address spruce growth promotion under well-watered conditions. To our knowledge, this is the first report of PGPB improving spruce growth under drought.

Overall, the 24-well system *in vivo* screening approach with its relatively high throughput possibility developed in this study turned out to be particularly useful as it prevented losing potential PGPB, which did not exhibit strong effects within the *in vitro* assays but showed their beneficial properties only when being applied to plants. At the same time, it verifies that potential PGP strains also exhibit their traits on the target plant and not only in an artificial assay or a model plant.

## 5. Conclusion

This study constitutes one of the first larger scale screenings for PGPM isolated from forest soils with axenic application to tree seedlings under drought stress. The double screening *in vitro* and *in planta* has proven beneficial in identifying PGP isolates that would have been overlooked based on the *in vitro* scoring alone. Overall, three microbes can be especially highlighted: *Streptomyces* Ke434 due to its high stress resistance, as well as *Caballeronia* Ke431 and *Paraburkholderia* Ke296 which showed positive effects on spruce growth under drought conditions. *Paraburkholderia* Ke296 also exhibited the most PGP traits of all tested strains. The *in planta* screening system offers a range of additional application possibilities, including adaptation to other plant species or testing further stress conditions such as heat, cold, and salt stress. Additionally, the system could be adapted for fungal inoculations or simultaneous inoculations together with bacteria.

## Data availability statement

The datasets generated or analyzed during the current study are included in this article and its supplementary information files or are available from the corresponding author upon reasonable request.

## Author contributions

Sonja Magosch: Conceptualization, Methodology, Validation, Formal analysis, Investigation, Data Curation, Visualization, Writing – Original Draft

Claudia Barrera: Methodology, Formal analysis, Investigation, Data Curation, Visualization, Writing – Review & Editing

Adrian Bölz: Formal analysis, Investigation

Karin Pritsch: Conceptualization, Supervision, Writing – Review & Editing

Michael Rothballer: Conceptualization, Funding acquisition, Supervision, Writing – Review & Editing

J. Philipp Benz: Conceptualization, Funding acquisition, Project administration, Supervision, Writing – Review & Editing

## Acknowledgments

We thank Fabian Weikl, Simin Rothballer and Petra Arnold for technical support and advice. Thank you to Theresa Kuhl-Nagel for providing the strains *Variovorax* sp. M92526_27 and *Luteibacter* sp. Cha2324a_16 as positive controls for the *in vitro* assays.

## Supplementary material

The Supplementary Material for this article can be found online.

