## Supplemental Tables and Figures for "Improving Seedling Survival for Forest Restorations: A Novel Screening Method to Identify Microbial Allies Against Drought Stress"

Sonja Magosch[1], Claudia Barrera[2], Adrian Bölz[1], Karin Pritsch[3], Michael Rothballer[4], J. Philipp Benz[1*]
[1]*Technical University of Munich, TUM School of Life Sciences, Professorship for Fungal Biotechnology in Wood Science, Freising, Germany*
[2]*Technical University of Munich, TUM School of Life Sciences, Professorship for Land Surface-Atmosphere Interactions, Freising, Germany*
[3] *Helmholtz Munich, Institute of Biochemical Plant Pathology, Neuherberg, Germany*
[4]*Helmholtz Munich, Institute of Network Biology, Neuherberg, Germany*

### Supplementary material

Table S1: Precipitation [mm] in the week before sampling at the two different sites.

| **Freising (Kranzberg)** | | **Kelheim** | |
|---|---|---|---|
| **Date** | **Precipitation [mm]** | **Date** | **Precipitation [mm]** |
| 07/21/2021 | 0 | 09/01/2021 | 0 |
| 07/22/2021 | 0 | 09/02/2021 | 0 |
| 07/23/2021 | 0 | 09/03/2021 | 0 |
| 07/24/2021 | 5 | 09/04/2021 | 0 |
| 07/25/2021 | 5.5 | 09/05/2021 | 0 |
| 07/26/2021 | 8.7 | 09/06/2021 | 0 |
| 07/27/2021 | 1 | 09/07/2021 | 0 |

Table S2: Bacterial strains utilized in this study including their isolation medium and origin.

| **Strain** | **Short name** | **Isolation medium** | **Origin** |
|---|---|---|---|
| *Pseudomonas* | Ke1 | M9 medium | Beech roots in Kelheim |
| *Paraburkholderia* | Ke15 | M9 medium | Beech roots in Kelheim |
| *Paraburkholderia* | Ke24 | M9 medium | Beech roots in Kelheim |
| *Rhizobium/Agrobacterium* | Ke26 | M9 medium | Beech roots in Kelheim |
| *Caballeronia* | Ke41 | M9 medium | Beech roots in Kelheim |
| *Caballeronia* | Ke57 | M9 medium | Spruce roots in Kelheim |
| *Bacillus* | Ke157 | King’s B medium | Beech roots in Kelheim |
| *Paraburkholderia* | Ke162 | King’s B medium | Beech roots in Kelheim |
| *Paraburkholderia* | Ke296 | MMNC medium | Beech roots in Kelheim |
| *Paraburkholderia* | Ke341 | MMNC medium | Beech roots in Kelheim |
| *Collimonas* | Ke374 | MMNC medium | Spruce roots in Kelheim |
| *Paraburkholderia* | Ke398 | MMNC medium | Spruce roots in Kelheim |
| *Caballeronia* | Ke431 | MMNC medium | Spruce roots in Kelheim |
| *Streptomyces* | Ke434 | KM4 agar | Beech roots in Kelheim |
| *Rhodococcus* | Ke442 | KM4 agar | Beech roots in Kelheim |
| *Streptomyces* | Ke462 | KM4 agar | Beech roots in Kelheim |
| *Rhodococcus* | Ke466 | KM4 agar | Beech roots in Kelheim |
| *Sporosarcina* | Ke477 | KM4 agar | Beech roots in Kelheim |
| *Streptomyces* | Ke502 | KM4 agar | Spruce roots in Kelheim |
| *Kitasatospora/Streptomyces* | Ke507 | KM4 agar | Spruce roots in Kelheim |
| *Bacillus* | KF3 | King’s B medium | Beech roots in Kranzberg |
| *Lysinibacillus* | KF19 | King’s B medium | Beech roots in Kranzberg |
| *Bacillus* | KF21 | King’s B medium | Beech roots in Kranzberg |
| *Bacillus* | KF47 | MMNC medium | Spruce roots in Kranzberg |
| *Pseudomonas* | KF80 | King’s B medium | Beech roots in Kranzberg |
| *Psychrobacillus* | KF94 | KM4 agar | Spruce roots in Kranzberg |
| *Viridibacillus* | KF108 | MMNC medium | Spruce roots in Kranzberg |
| *Herbaspirillum* | KF115 | MMNC medium | Spruce roots in Kranzberg |
| *Caballeronia* | KF127 | MMNC medium | Beech roots in Kranzberg |
| *Streptomyces* | KF143 | King’s B medium | Beech roots in Kranzberg |
| *Streptomyces* | KF147 | MMNC medium | Spruce roots in Kranzberg |
| *Pseudomonas* | KF164 | M9 medium | Beech roots in Kranzberg |
| *Streptomyces* | KF207 | KM4 agar | Spruce roots in Kranzberg |
| *Streptomyces* | KF209 | MMNC medium | Spruce roots in Kranzberg |
| *Streptomyces* | KF215 | KM4 agar | Spruce roots in Kranzberg |
| *Planktothrix/Variovorax* | KF227 | M9 medium | Spruce roots in Kranzberg |

Table S3: Fungal strains utilized in this study including their isolation medium and origin.

| Strain | Short name | Isolation medium | Origin |
|---|---|---|---|
| *Umbelopsis* | F4 | MMNC medium | Beech roots in Kelheim |
| *Tolypocladium* | F5 | MMNC medium | Spruce roots in Kelheim |
| *Metapochonia/Pochonia* | F6 | M9 medium | Spruce roots in Kelheim |
| *Umbelopsis* | F10 | MMNC medium | Spruce roots in Kelheim |
| *Umbelopsis* | F11 | MMNC medium | Spruce roots in Kelheim |
| *Umbelopsis* | F12 | MMNC medium | Spruce roots in Kelheim |
| *Umbelopsis* | F14 | MMNC medium | Spruce roots in Kelheim |
| *Lycoperdon* | F31 | MMNC medium | Fruiting body near beech roots in Kelheim |
| *Gymnopus* | F32 | MMNC medium | Fruiting body near spruce roots in Kelheim |
| *Umbelopsis* | F34 | MMNC medium | Spruce roots in Kelheim |
| *Tolypocladium* | F36 | MMNC medium | Spruce roots in Kelheim |
| *Podila/uncultured soil fungus* | F37 | MMNC medium | Spruce roots in Kelheim |
| *Mortierella/Podila* | F40 | King’s B medium | Spruce roots in Kelheim |
| *Umbelopsis* | F7 | MMNC medium | Beech roots in Kranzberg |
| *Umbelopsis* | F17 | MMNC medium | Beech roots in Kranzberg |
| *Mortierella* | F38 | MMNC medium | Beech roots in Kranzberg |
| *Mortierella* | F39 | MMNC medium | Spruce roots in Kranzberg |

Table S4: PEG, NaCl and $H^+$ concentrations applied during the stress tolerance assays, and scoring.

| PEG [MPa] | NaCl [%] | pH ($H^+$) | Scoring |
|---|---|---|---|
| 0 | 0 | 7 and 8 | 0 |
| -0.25 | 0.1 | 6 | 1 |
| -0.5 | 2.5 | - | 1 |
| -0.75 | 3.5 | 5 | 2 |
| -1.0 | 5.5 | - | 2 |
| -1.25 | 7.5 | 4 | 3 |
| -1.5 | 12 | 3 | 3 |
| -1.75 | 15 | 2 | 3 |

Table S5: *In vitro* assays of the utilized positive controls. “-“ = not tested.

| Strain name | ACC deaminase production | | | N-fixation | | | P-solubilization |
|---|---|---|---|---|---|---|---|
| | M9 | M9+ ACC | M9 w/o ACC | Nfb agar | | Jensen's agar | |
| *Variovorax* M92526_27 | 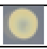 | 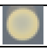 | 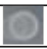 | - | - | - | - |
| *Luteibacter* Cha23&24_a1 | - | - | - | - | - | - | 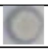 |
| *E. coli* DH5alpha (MGC3) | - | - | - | 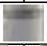 | | 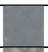 | - |
| *Azospirillum brasilense* Sp7 | - | - | - | 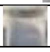 | 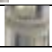 | 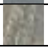 | - |

Table S6: Recovery of bacteria from the 2-3 highest NaCl, PEG and $H^+$ concentrations. The highest concentration from which strains could recover is indicated. Non-recovery of strains is indicated by "-".

| Strain name | NaCl | PEG | $H^+$ |
|---|---|---|---|
| *Pseudomonas* Ke1 | - | -1.25 | - |
| *Paraburkholderia* Ke15 | - | -1.5 | 4 |
| *Paraburkholderia* Ke24 | - | - | 4 |
| *Rhizobium* Ke26 | - | - | 4 |
| *Caballeronia* Ke41 | - | - | 4 |
| *Caballeronia* Ke57 | 15 | -1.75 | 3 |
| *Bacillus* Ke157 | 15 | -1.75 | 2 |
| *Paraburkholderia* Ke162 | - | - | 4 |
| *Paraburkholderia* Ke296 | 15 | -1.25 | 4 |
| *Paraburkholderia* Ke341 | 15 | -1.75 | 4 |
| *Collimonas* Ke374 | 12 | -1.75 | 4 |
| *Paraburkholderia* Ke398 | 15 | -1.75 | 4 |
| *Caballeronia* Ke431 | - | - | 4 |
| *Streptomyces* Ke434 | 15 | -1.75 | 3 |
| *Rhodococcus* Ke442 | 15 | -1.75 | 3 |
| *Streptomyces* Ke462 | 15 | -1.75 | 4 |
| *Rhodococcus* Ke466 | 15 | -1.5 | 2 |
| *Sporosarcina* Ke477 | 15 | -1.75 | - |
| *Streptomyces* Ke502 | 15 | -1.75 | 3 |
| *Streptomyces/Kitasatospora* Ke507 | 15 | -1.75 | 3 |
| *Bacillus* KF3 | 15 | -1.75 | 3 |
| *Lysinibacillus* KF19 | 15 | -1.75 | - |
| *Bacillus* KF21 | 15 | -1.75 | 4 |
| *Bacillus* KF47 | 15 | -1.75 | 3 |
| *Pseudomonas* KF80 | - | - | - |
| *Psychrobacillus* KF94 | - | -1.75 | 4 |
| *Viridibacillus* KF108 | 15 | -1.75 | 2 |
| *Herbaspirillum* KF115 | - | - | - |
| *Caballeronia* KF127 | - | - | - |
| *Streptomyces* KF143 | 15 | -1.75 | - |
| *Streptomyces* KF147 | 15 | -1.75 | - |
| *Pseudomonas* KF164 | - | - | - |
| *Streptomyces* KF207 | 12 | -1.75 | 4 |
| *Streptomyces* KF209 | 12 | -1.75 | 4 |
| *Streptomyces* KF215 | 15 | -1.75 | - |
| *Planktothrix/Variovorax* KF227 | - | - | - |

Table S7: Bacterial IAA production with and without tryptophane in µg $mL^{-1}$ with the respective standard deviation. PC = positive control.

| Strain name | IAA production [µg $mL^{-1}$] | |
|---|---|---|
| | With tryptophane | Without tryptophane |
| *Pseudomonas* Ke1 | 25.4 ± 7.0 | 6.5 ± 0.3 |
| *Paraburkholderia* Ke15 | 7.8 ± 1.9 | 6.2 ± 2.7 |
| *Paraburkholderia* Ke24 | 9.8 ± 3.8 | 8.2 ± 5.6 |
| *Rhizobium* Ke26 | 10.6 ± 2.1 | 9.9 ± 0.8 |
| *Caballeronia* Ke41 | 8.3 ± 5.9 | 2.3 ± 1.5 |
| *Caballeronia* Ke57 | 10.4 ± 1.0 | 9.2 ± 6.8 |
| *Bacillus* Ke157 | 10.9 ± 1.0 | 1.1 ± 0.8 |
| *Paraburkholderia* Ke162 | 5.7 ± 1.4 | 6.0 ± 3.4 |
| *Paraburkholderia* Ke296 | 2.7 ± 1.2 | 1.8 ± 1.2 |
| *Paraburkholderia* Ke341 | 7.7 ± 2.9 | 9.6 ± 7.0 |
| *Collimonas* Ke374 | 10.1 ± 3.0 | 5.3 ± 2.2 |
| *Paraburkholderia* Ke398 | 2.3 ± 1.2 | 1.4 ± 0.4 |
| *Caballeronia* Ke431 | 10.0 ± 1.0 | 5.8 ± 3.2 |
| *Streptomyces* Ke434 | 1.4 ± 0.6 | 0.4 ± 0.1 |
| *Rhodococcus* Ke442 | 0.5 ± 0.4 | 0.4 ± 0.6 |
| *Streptomyces* Ke462 | 7.7 ± 0.7 | 3.6 ± 2.4 |
| *Rhodococcus* Ke466 | 1.9 ± 0.2 | 0.1 ± 0.1 |

| | | |
|---|---|---|
| *Sporosarcina* Ke477 | 18.0 ± 7.8 | 6.6 ± 3.0 |
| *Streptomyces* Ke502 | 2.2 ± 0.3 | 3.2 ± 0.2 |
| *Streptomyces/Kitasatospora* Ke507 | 1.2 ± 0.0 | 0.3 ± 0.1 |
| *Bacillus* KF3 | 11.0 ± 3.1 | 2.3 ± 0.6 |
| *Lysinibacillus* KF19 | 41.1 ± 12.3 | 2.7 ± 1.5 |
| *Bacillus* KF21 | 6.1 ± 1.1 | 1.6 ± 0.5 |
| *Bacillus* KF47 | 8.6 ± 3.5 | 2.3 ± 1.1 |
| *Pseudomonas* KF80 | 24.4 ± 4.6 | 8.4 ± 1.4 |
| *Psychrobacillus* KF94 | 19.9 ± 11.6 | 10.0 ± 12.6 |
| *Viridibacillus* KF108 | 55.7 ± 11.8 | 14.2 ± 9.2 |
| *Herbaspirillum* KF115 | 11.0 ± 2.1 | 4.6 ± 1.1 |
| *Caballeronia* KF127 | 39.4 ± 2.7 | 7.5 ± 3.1 |
| *Streptomyces* KF143 | 11.6 ± 1.9 | 2.8 ± 2.4 |
| *Streptomyces* KF147 | 19.9 ± 12.1 | 8.4 ± 13.6 |
| *Pseudomonas* KF164 | 39.4 ± 3.4 | 4.9 ± 1.8 |
| *Streptomyces* KF207 | 20.7 ± 3.0 | 20.3 ± 2.4 |
| *Streptomyces* KF209 | 1.1 ± 0.7 | 1.6 ± 0.9 |
| *Streptomyces* KF215 | 16.6 ± 8.7 | 7.0 ± 8.7 |
| *Planktothrix/Variovorax* KF227 | 1.0 ± 0.1 | 1.7 ± 0.2 |
| *Herbaspirillum frisingense* GSF30 (PC) | 12.4 ± 2.1 | 8.5 ± 1.0 |

Table S8: Fungal IAA production with and without tryptophane in µg $mg^{-1}$ with the respective standard deviation.

| **Strain name** | **IAA production [µg $mg^{-1}$]** | |
|---|---|---|
| | **With tryptophane** | **Without tryptophane** |
| *Umbelopsis* F4 | 0.36 ± 0.15 | 0.04 ± 0.02 |
| *Tolypocladium* F5 | 0.50 ± 0.20 | 0.02 ± 0.02 |
| *Metapochonia*/*Pochonia* F6 | 0.05 ± 0.03 | 0.02 ± 0.03 |
| *Umbelopsis* F7 | 0.59 ± 0.38 | 0.03 ± 0.02 |
| *Umbelopsis* F10 | 0.89 ± 0.08 | 0.02 ± 0.00 |
| *Umbelopsis* F11 | 5.53 ± 2.25 | 0.13 ± 0.05 |
| *Umbelopsis* F12 | 0.43 ± 0.15 | 0.00 ± 0.01 |
| *Umbelopsis* F14 | 1.75 ± 0.42 | 0.01 ± 0.00 |
| *Umbelopsis* F17 | 0.82 ± 0.34 | -0.02 ± 0.06 |
| Fruiting body, *Lycoperdon* F31 | 1.74 ± 0.09 | 0.00 ± 0.01 |
| Fruiting body, *Gymnopus* F32 | 1.78 ± 1.07 | -0.05 ± 0.01 |
| *Umbelopsis* F34 | 0.90 ± 0.27 | -0.02 ± 0.02 |
| *Tolypocladium* F36 | 1.41 ± 0.23 | 0.04 ± 0.04 |
| *Podila*/uncultured soil fungus F37 | 2.21 ± 0.37 | 0.01 ± 0.01 |
| *Mortierella* F38 | 4.73 ± 1.71 | -0.02 ± 0.04 |
| *Mortierella* F39 | 6.35 ± 1.46 | -0.08 ± 0.06 |
| *Mortierella*/*Podila* F40 | 3.99 ± 1.29 | -0.03 ± 0.03 |

Table S9: Summary of the effects of the isolates on plant growth promotion in comparison their respective control in the 24-well test system. Significant plant growth promotion after inoculation is marked with “+”. In the experiment with Ke431 and Ke434 inoculation, dry weight measurement was not performed for the control plants (n.c.). Bacteria are listed in descending order of importance with respect to their ability to promote plant growth.

| **Inoculated bacteria** | **Significant plant growth promotion, indicated by “+”** | | | | | | | | | | | | | |
|---|---|---|---|---|---|---|---|---|---|---|---|---|---|---|
| **Strain name** | **Seedling length** | | **Seedling fresh weight** | | **Root length** | | **Root fresh weight** | | **Shoot length** | | **Shoot fresh weight** | | **Seedling dry weight** | |
| | **WW** | **DS** | **WW** | **DS** | **WW** | **DS** | **WW** | **DS** | **WW** | **DS** | **WW** | **DS** | **WW** | **DS** |
| *Caballeronia* Ke431 | **+** | **+** | **+** | | **+** | **+** | **+** | | | | **+** | | n.c. | |
| *Paraburkholderia* Ke296 | | **+** | | | | **+** | | | | | | | | |
| *Paraburkholderia* Ke15 | **+** | | **+** | | **+** | | **+** | | **+** | | **+** | | **+** | |
| *Paraburkholderia* Ke24 | **+** | | **+** | | **+** | | **+** | | | | **+** | | **+** | |
| *Herbaspirillum* KF115 | | | **+** | | | | **+** | | | | **+** | | **+** | |
| *Streptomyces* KF215 | **+** | | | | **+** | | **+** | | | | | | | |
| *Paraburkholderia* Ke162 | | | | | | | **+** | | | | | | | |
| *Psychrobacillus* KF94 | | | | | | | | | | | | | | **+** |
| *Rhodococcus* Ke466 | | | | | | | | | | | | | **+** | |
| *Bacillus* KF21 | | | | | | | | | | | | | **+** | |
| No effect *in vivo: Pseudomonas* Ke1, *Caballeronia* Ke41, *Caballeronia* Ke57, *Paraburkholderia* Ke431, *Collimonas* Ke374, *Paraburkholderia* Ke398, *Streptomyces* Ke434, *Rhodococcus* Ke442, *Streptomyces* Ke462, *Sporosarcina* Ke477, *Streptomyces/Kitasatospora* Ke507, *Bacillus* KF3, *Bacillus* KF47, *Viridibacillus* KF108, *Caballeronia* KF127, *Streptomyces* KF143, *Streptomyces* KF147, *Pseudomonas* KF164, *Streptomyces* KF207 | | | | | | | | | | | | | | |

Table S10: Summary of the plant survival rates and colony forming units (CFUs) after inoculation with the different isolates and growth for 3 weeks in a phytochamber. Survival rates are indicated in relation to the respective uninoculated control plants of each experiment as relative values. An improvement of the survival rate is highlighted in grey.

| Short name | Genus | Increase/decrease in survival (fold change, relative values) | | CFU count | |
|---|---|---|---|---|---|
| | | WW | DS | WW | DS |
| Ke1 | *Pseudomonas* | 1.47 | 2.0 | $1.5 \times 10^6$ | $2.6 \times 10^5$ |
| Ke15 | *Paraburkholderia* | 1.35 | 0.75 | $3.1 \times 10^7$ | $2.5 \times 10^7$ |
| Ke24 | *Paraburkholderia* | 1.18 | 0.75 | $4.9 \times 10^7$ | $1.5 \times 10^7$ |
| Ke41 | *Caballeronia* | 0.8 | 0.89 | $1.5 \times 10^7$ | $7.5 \times 10^6$ |
| Ke57 | *Caballeronia* | 0.94 | 0.71 | $6.8 \times 10^6$ | $3.4 \times 10^5$ |
| Ke162 | *Paraburkholderia* | 0.91 | 0.72 | $1.3 \times 10^7$ | $1.8 \times 10^7$ |
| Ke296 | *Paraburkholderia* | 1.11 | 1.18 | $5.7 \times 10^6$ | $4.8 \times 10^6$ |
| Ke341 | *Paraburkholderia* | 0.83 | 0.87 | $3.9 \times 10^7$ | $3.5 \times 10^7$ |
| Ke374 | *Collimonas* | 1.0 | 1.07 | $3.0 \times 10^7$ | $1.0 \times 10^7$ |
| Ke398 | *Paraburkholderia* | 0.83 | 0.63 | $2.7 \times 10^7$ | $7.7 \times 10^7$ |
| Ke431 | *Caballeronia* | 1.0 | 3.01 | $6.2 \times 10^5$ | $2.3 \times 10^5$ |
| Ke434 | *Streptomyces* | 1.11 | 1.0 | $1.8 \times 10^6$ | $5.0 \times 10^6$ |
| Ke442 | *Rhodococcus* | 0.96 | 0.72 | $1.3 \times 10^6$ | $1.8 \times 10^6$ |
| Ke462 | *Streptomyces* | 0.83 | 1.0 | $1.6 \times 10^4$ | $8.7 \times 10^5$ |
| Ke466 | *Rhodococcus* | 0.94 | 0.5 | $2.1 \times 10^6$ | $4.0 \times 10^6$ |
| Ke477 | *Sporosarcina* | 0.72 | 1.36 | $2.8 \times 10^6$ | $1.2 \times 10^4$ |
| Ke507 | *Streptomyces/Kitasatospora* | 0.61 | 0 | $2.5 \times 10^5$ | dead |
| KF3 | *Bacillus* | 1.1 | 2.0 | $4.8 \times 10^6$ | $1.6 \times 10^4$ |
| KF21 | *Bacillus* | 1.0 | 0.25 | $9.3 \times 10^4$ | $7.4 \times 10^5$ |
| KF47 | *Bacillus* | 1.1 | 1.78 | $2.5 \times 10^4$ | $1.4 \times 10^4$ |
| KF94 | *Psychrobacillus* | 1.2 | 2.83 | - | - |
| KF108 | *Viridibacillus* | 1.47 | 3.16 | - | $4.9 \times 10^3$ |
| KF115 | *Herbaspirillum* | 1.17 | 1.57 | $1.5 \times 10^7$ | $2.9 \times 10^6$ |
| KF127 | *Caballeronia* | 0.78 | 1.36 | $3.6 \times 10^7$ | $7.0 \times 10^6$ |
| KF143 | *Streptomyces* | 0.83 | 0.63 | $6.6 \times 10^6$ | $4.8 \times 10^6$ |
| KF147 | *Streptomyces* | 1.47 | 1.0 | $8.9 \times 10^5$ | $3.1 \times 10^6$ |
| KF164 | *Pseudomonas* | 0.91 | 1.14 | $1.3 \times 10^6$ | $9.6 \times 10^5$ |
| KF207 | *Streptomyces* | 0.8 | 0.67 | $1.5 \times 10^5$ | $3.6 \times 10^5$ |
| KF215 | *Streptomyces* | 1.0 | 1.18 | $4.0 \times 10^5$ | $1.9 \times 10^6$ |

Table S11: Summary of the PGP properties and stress tolerance of the seven PGPB identified within the 24-well-based test system. The assigned scores for PEG, NaCl, and pH tolerance, as well as for IAA-production, are indicated. ACC-utilization, N-fixation, P-solubilization and siderophore (S) production is marked with “X”.

| Short | Genus | PEG | NaCl | pH | IAA | ACC | N | P | S |
|---|---|---|---|---|---|---|---|---|---|
| Ke15 | *Paraburkholderia* | 2 | 1 | 3 | 1 | X | - | X | - |
| Ke24 | *Paraburkholderia* | 2 | 1 | 3 | 1 | X | - | X | - |
| Ke162 | *Paraburkholderia* | 2 | 1 | 3 | 1 | X | - | X | X |
| Ke296 | *Paraburkholderia* | 2 | 1 | 3 | - | X | X | X | - |
| Ke431 | *Caballeronia* | 1 | 1 | 2 | 2 | X | X | - | - |
| KF115 | *Herbaspirillum* | 1 | 1 | 2 | 2 | X | - | - | X |
| KF215 | *Streptomyces* | 3 | 2 | 2 | 3 | - | - | - | - |

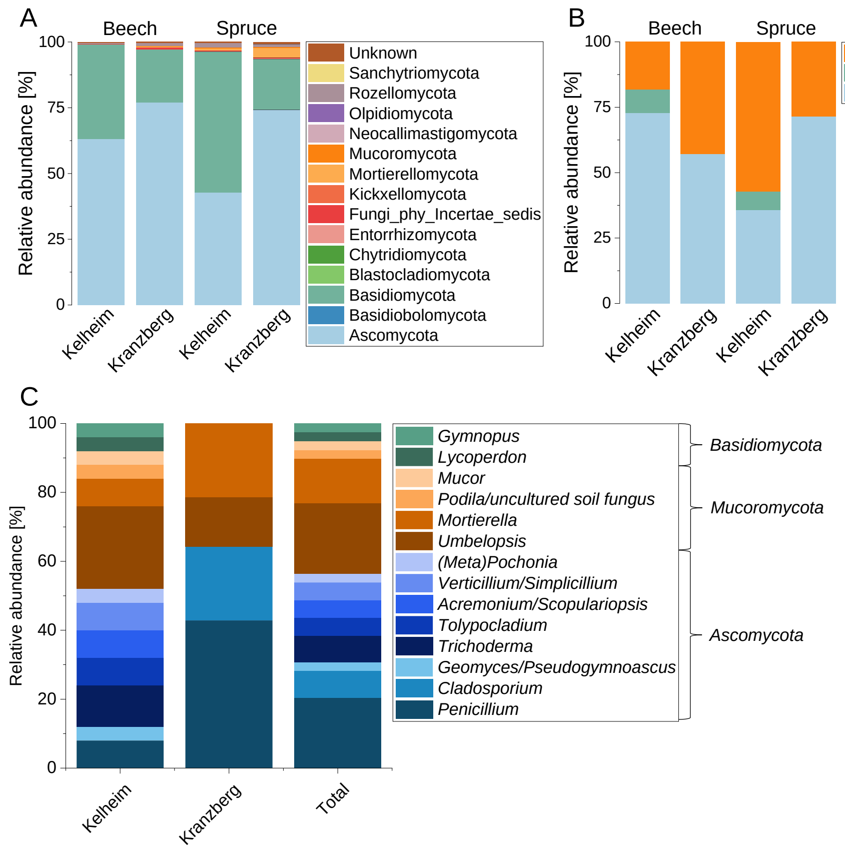


Figure S1: Comparison of the relative abundance of fungal phyla in rhizosphere communities and root isolates of beech and spruce in Kelheim and Kranzberg. (A) Relative abundance [%] of the fungal phyla in beech and spruce rhizosphere communities in Kelheim and Kranzberg. (B) Relative abundance [%] of the fungal phyla isolated as single strains from beech and spruce roots in Kelheim and Kranzberg. (C) Relative abundance [%] of the fungal genera isolated as single strains in Kelheim and Kranzberg, and at both sites combined. Fungi of the same phyla are shown in similar colors. Ascomycota: blue. Basidiomycota: green. Mucoromycota: brown. The total number of isolates (n) is given below each diagram. More closely related genera are depicted in similar colors.

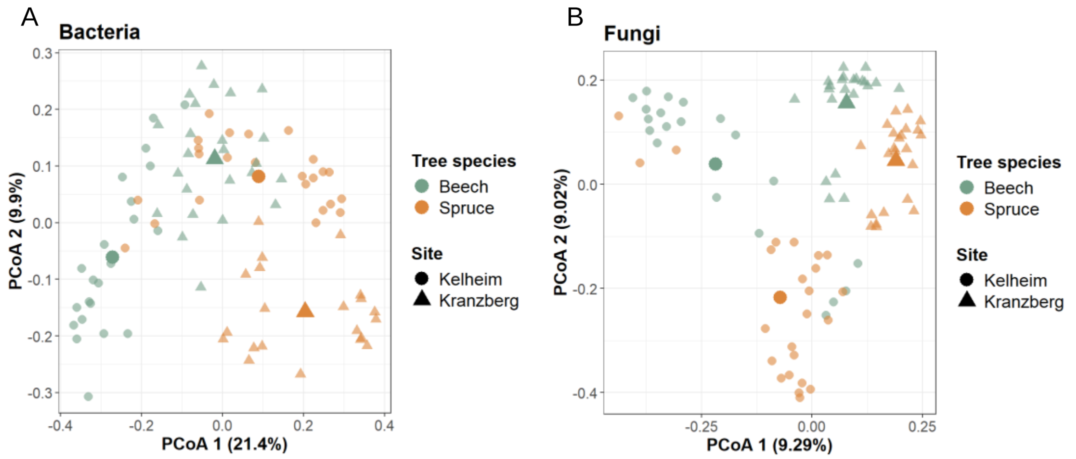


Figure S2: Principal Coordinate Analysis (PCoA) showing the site- and tree species-related differences in the (A) bacterial and (B) fungal rhizosphere community composition. The average of each combination is highlighted by a more intense color and a bigger size of the respective symbols.

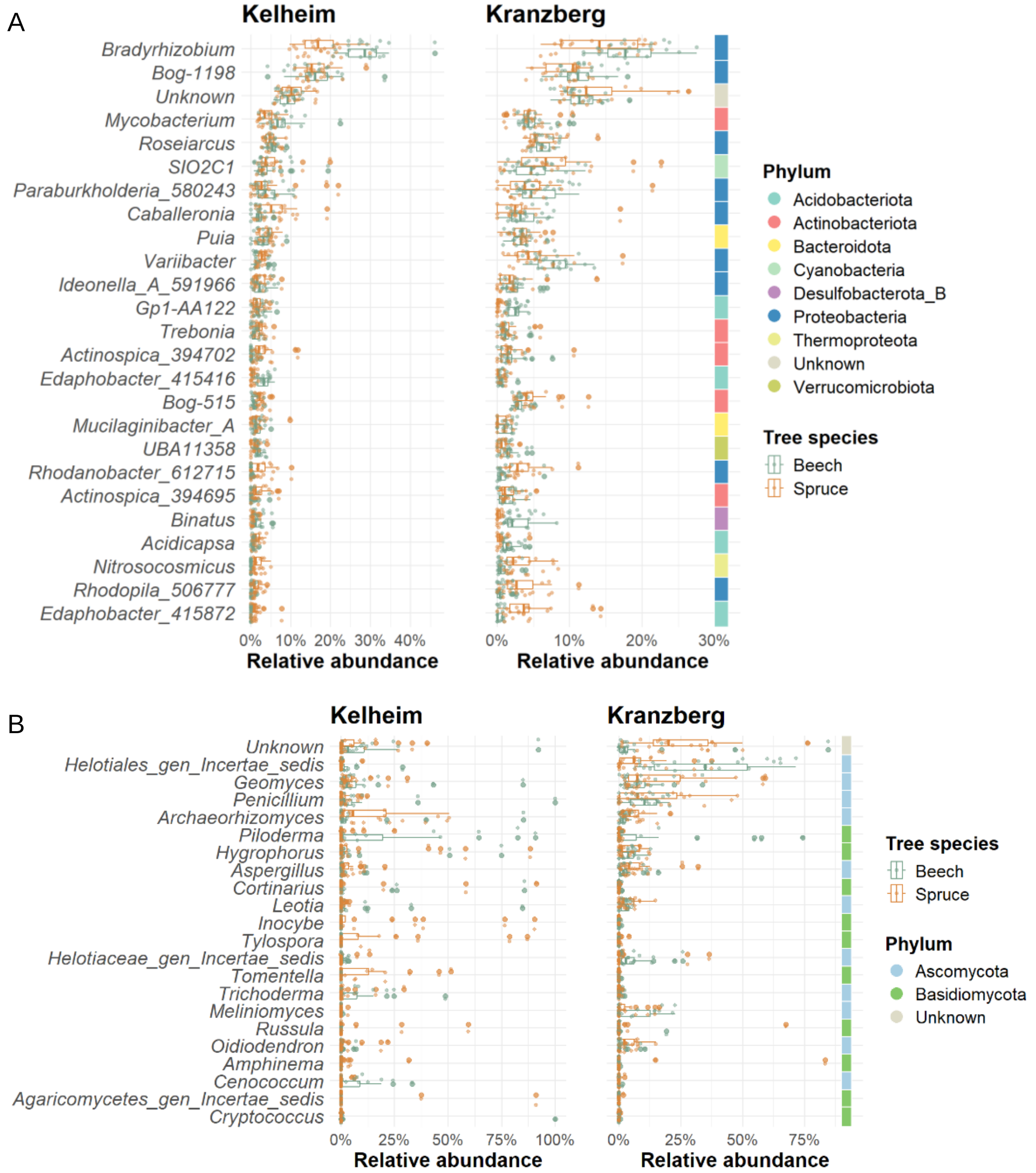


Figure S3: Relative abundance [%] of the (A) bacterial and (B) fungal genera in beech and spruce rhizosphere communities in Kelheim and Kranzberg, sorted according to their global abundance (descending order). The colors on the right indicate their phylum affiliation.

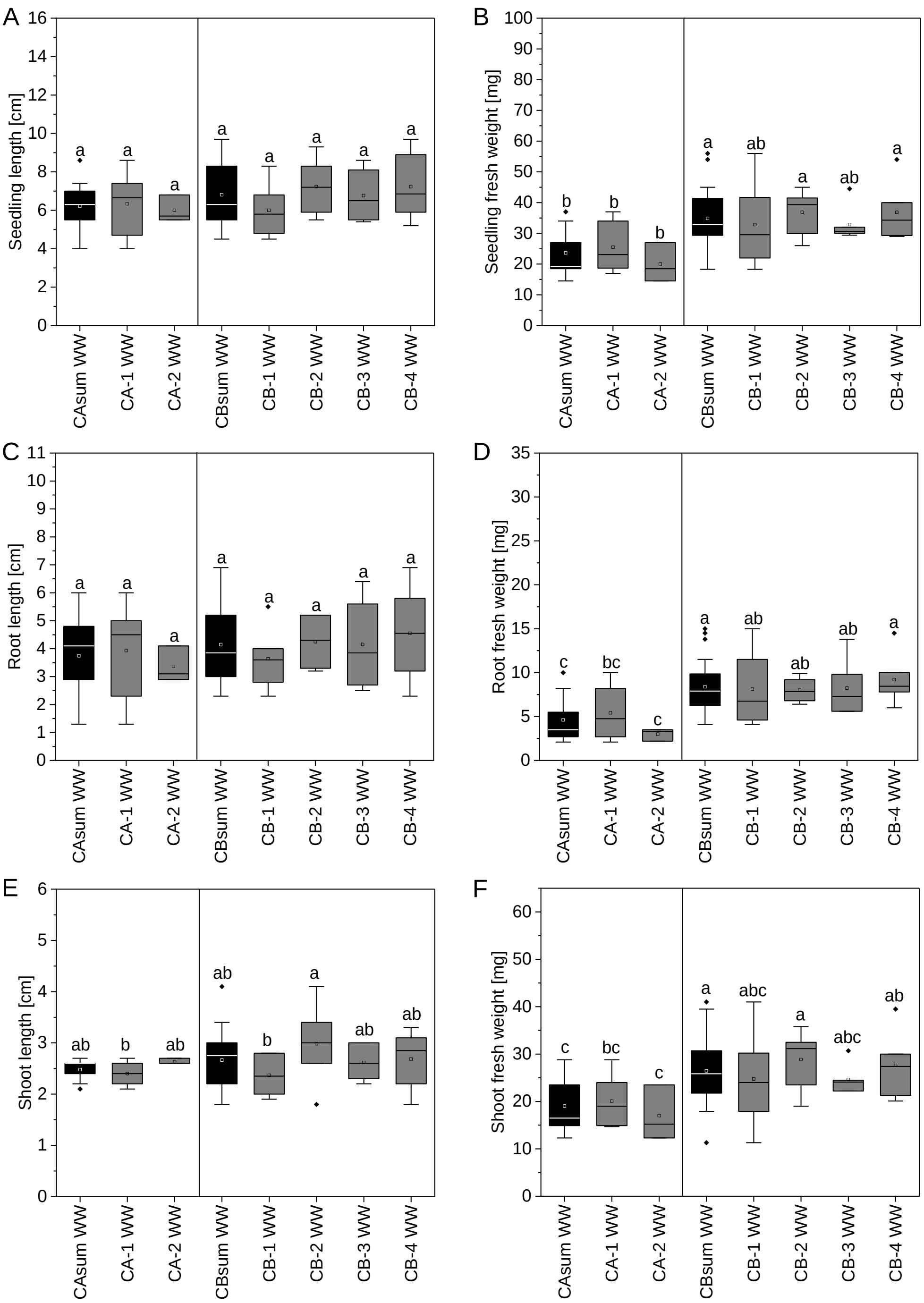


Figure S4: Comparison of the initially used internal controls under well-watered conditions. CAsum is the average of the internal controls CA1 and CA2. CBsum is the average of the internal controls CB1-CB4. There is no significant difference between the average of the internal controls and the internal controls.

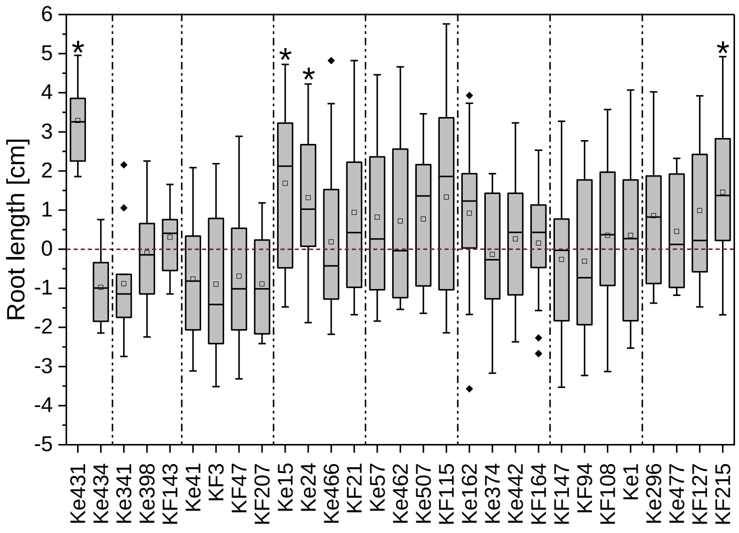


Figure S5: Increase or decrease in root length under well-watered conditions after inoculation with the bacterial isolates in comparison to the control plants of each experiment (red zero baseline). The single experiments are separated by dashed lines. Each positive value indicates an absolute increase in root length of the inoculated plants compared to the respective controls, while negative values mark a decrease. Significant plant growth-promotion after inoculation with the isolates compared to the respective control is marked with asterisks.

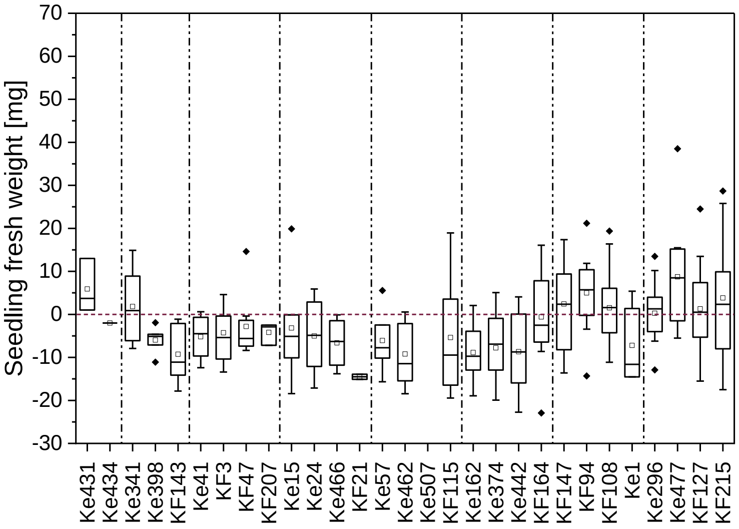


Figure S6: Increase or decrease in seedling fresh weight under well-watered conditions after inoculation with the bacterial isolates in comparison to the control plants of each experiment (red zero baseline). The single experiments are separated by dashed lines. Each positive value indicates an absolute increase in root length of the inoculated plants compared to the respective controls, while negative values mark a decrease. Significant plant growth-promotion after inoculation with the isolates compared to the respective control is marked with asterisks.

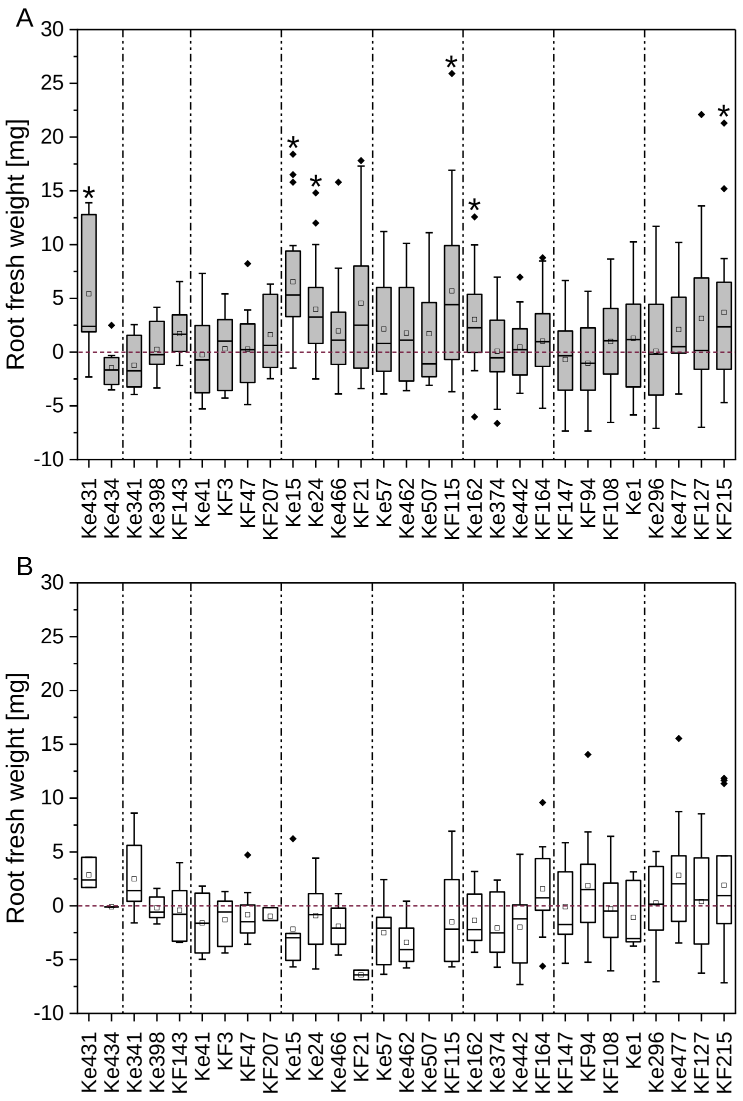


Figure S7: Increase or decrease in root fresh weight under (A) well-watered and (B) drought stress conditions after inoculation with the bacterial isolates in comparison to the control plants of each experiment (red zero baseline). The single experiments are separated by dashed lines. Each positive value indicates an absolute increase in root fresh weight of the inoculated plants compared to the respective controls, while negative values mark a decrease. Significant plant growth-promotion after inoculation with the isolates compared to the respective control is marked with asterisks.

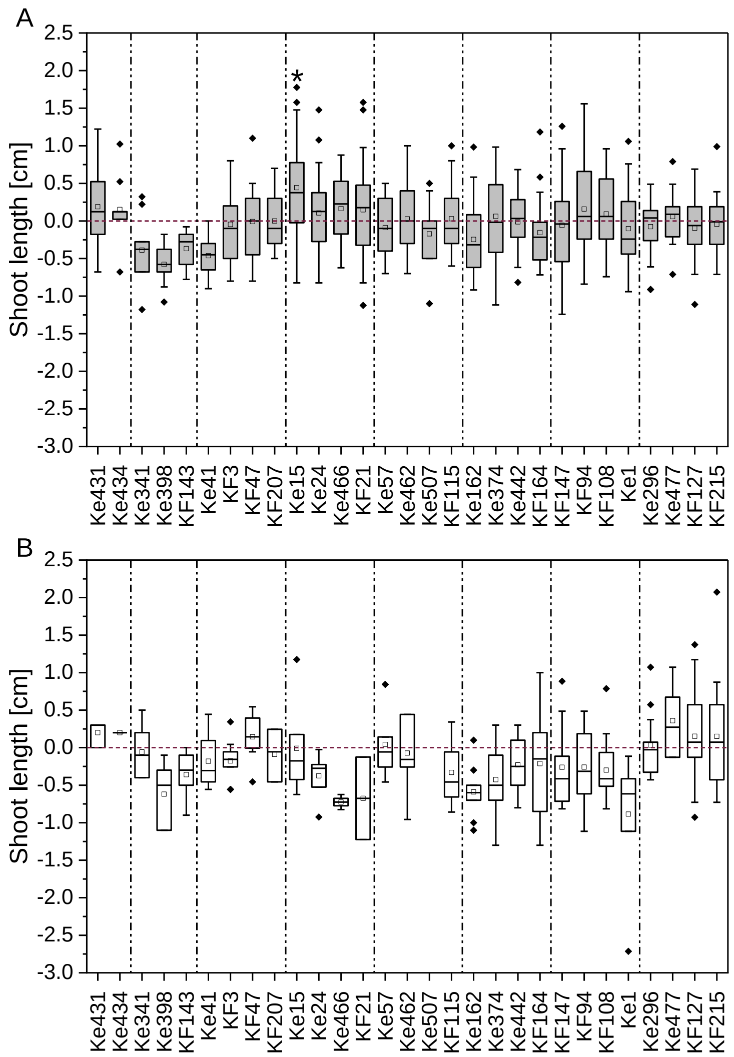


Figure S8: Increase or decrease in shoot length under (A) well-watered and (B) drought stress conditions after inoculation with the bacterial isolates in comparison to the control plants of each experiment (red zero baseline). The single experiments are separated by dashed lines. Each positive value indicates an absolute increase in shoot length of the inoculated plants compared to the respective controls, while negative values mark a decrease. Significant plant growth-promotion after inoculation with the isolates compared to the respective control is marked with asterisks.

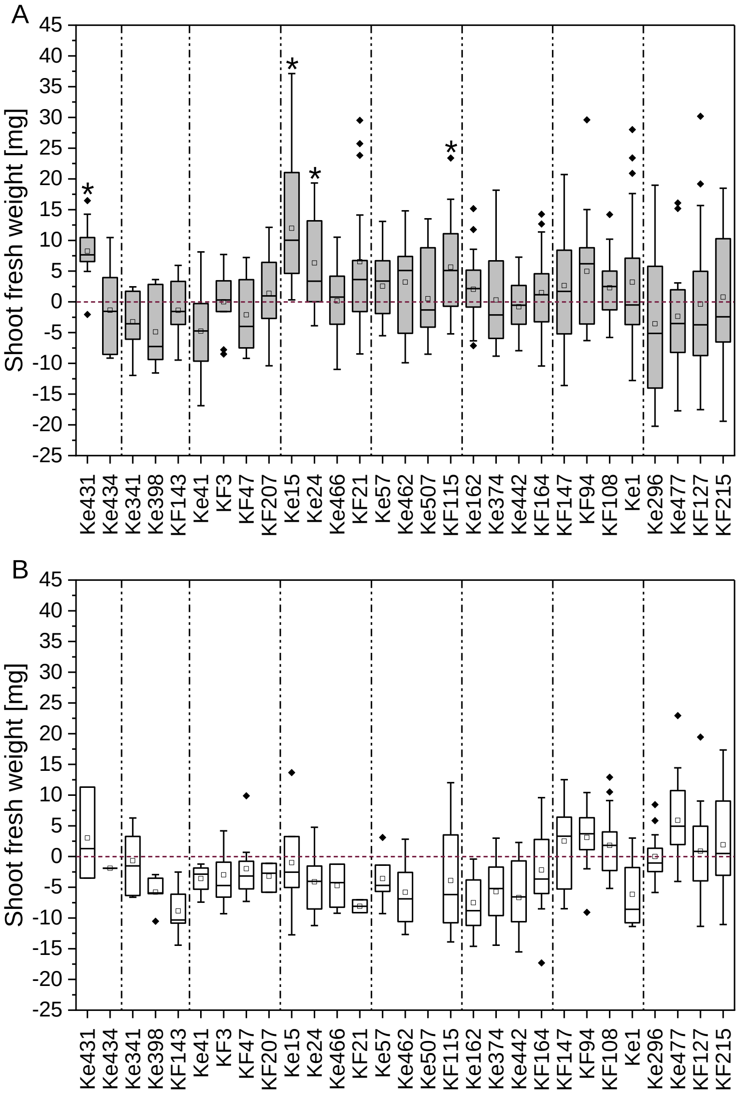


Figure S9: Increase or decrease in shoot fresh weight under (A) well-watered and (B) drought stress conditions after inoculation with the bacterial isolates in comparison to the control plants of each experiment (red zero baseline). The single experiments are separated by dashed lines. Each positive value indicates an absolute increase in shoot fresh weight of the inoculated plants compared to the respective controls, while negative values mark a decrease. Significant plant growth-promotion after inoculation with the isolates compared to the respective control is marked with asterisks.

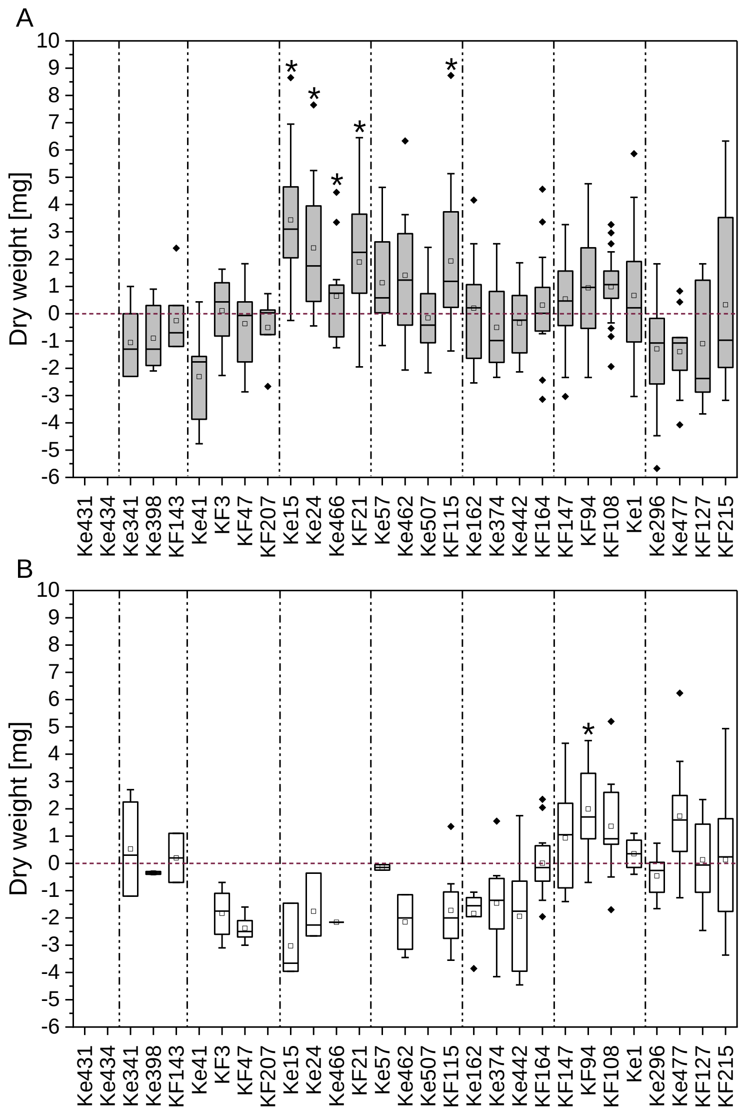


C

| Short name | Number of plants available for determination of dry weight | |
|---|---|---|
| | WW | DS |
| Ke1 | 20 (14) | 4 (4) |
| Ke15 | 20 (14) | 3 (5) |
| Ke24 | 17 (14) | 3 (5) |
| Ke41 | 5 (12) | 0 (2) |
| Ke57 | 14 (15) | 2 (4) |
| Ke162 | 18 (20) | 7 (11) |
| Ke296 | 17 (15) | 10 (8) |
| Ke341 | 7 (16) | 4 (8) |
| Ke374 | 20 (20) | 12 (11) |
| Ke398 | 7 (16) | 2 (8) |
| Ke431 | n.t. | n.t. |
| Ke434 | n.t. | n.t. |
| Ke442 | 19 (20) | 7 (11) |
| Ke462 | 12 (15) | 4 (4) |
| Ke466 | 13 (14) | 1 (5) |
| Ke477 | 10 (15) | 12 (8) |
| Ke507 | 8 (15) | 0 (4) |
| KF3 | 8 (12) | 6 (2) |
| KF21 | 14 (14) | 0 (5) |
| KF47 | 9 (12) | 5 (2) |
| KF94 | 16 (14) | 15 (4) |
| KF108 | 7 (16) | 6 (8) |
| KF115 | 18 (15) | 8 (4) |
| KF127 | 11 (15) | 12 (8) |
| KF143 | 7 (16) | 2 (8) |
| KF147 | 20 (14) | 10 (4) |
| KF164 | 18 (20) | 13 (11) |
| KF207 | 5 (12) | 0 (2) |
| KF215 | 15 (15) | 11 (8) |

Figure S10: Increase or decrease in seedling dry weight under (A) well-watered (WW) and (B) drought stress (DS) conditions after inoculation with the bacterial isolates in comparison to the control plants of each experiment (red zero baseline). The single experiments are separated by dashed lines. Each positive value indicates an absolute increase in dry weight of the inoculated plants compared to the respective controls, while negative values mark a decrease. Significant plant growth-promotion after inoculation with the isolates compared to the respective control is marked with asterisks. (C) The number of plants available for dry weight determination depended on the survival rate and excluded plants used for CFU count. The number of control plants is given in brackets.

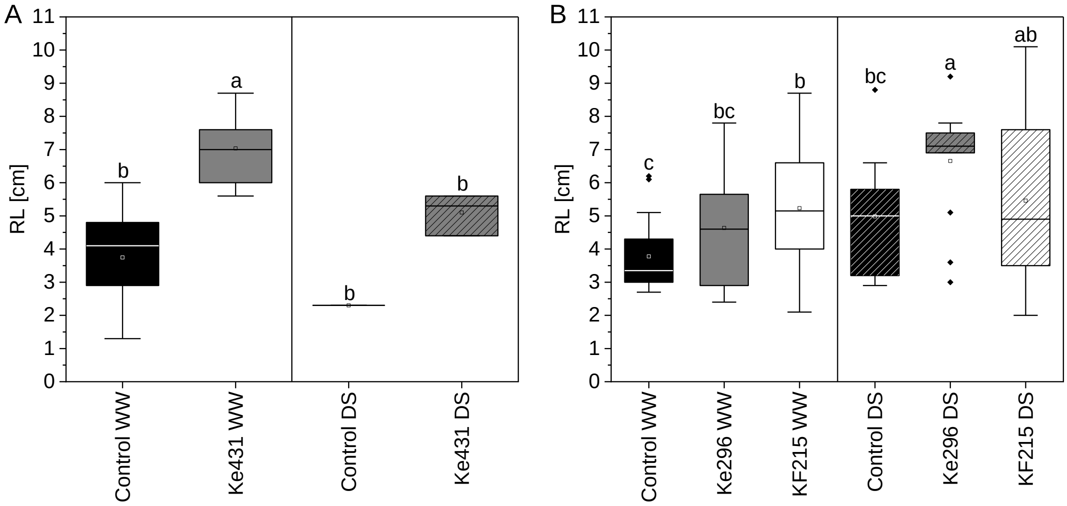


Figure S11: Root length after inoculation with (A) Ke431 and (B) Ke296 and KF215.
